# Impact of genus (*Geotrigona*, *Melipona*, *Scaptotrigona*) in the ^1^H-NMR organic profile and authenticity test of honey processed in cerumen pots by stingless bees in Ecuador

**DOI:** 10.1101/2022.05.17.492391

**Authors:** Patricia Vit, Jane van der Meulen, Silvia RM Pedro, Isabelle Esperança, Rahimah Zakaria, Gudrun Beckh, Favian Maza

**Affiliations:** Apitherapy and Bioactivity, Food Science Department, Faculty of Pharmacy and Bioanalysis, Universidad de Los Andes, Mérida 5101, Venezuela; Quality Services International GmbH, 28199 Bremen, Germany; Biology Department, Universidade de São Paulo, Ribeirão Preto, Brazil; Institute of Chemistry, Universidad Federal de Rio de Janeiro, Rio de Janeiro, RJ 21945970, Brazil; Department of Physiology, School of Medical Sciences, Universiti Sains Malaysia, Kota Bharu, Malaysia; Faculty of Agricultural and Livestock Sciences, Universidad Técnica de Machala, Machala, El Oro province, Ecuador

**Keywords:** *Geotrigona*, HATIE, HBT, *Melipona*, Meliponini, ^1^H-NMR, pot-honey, *Scaptotrigona*, stingless bees

## Abstract

The biodiversity of Ecuadorian stingless bees is almost 200 species. Traditional pot-honey harvest in Ecuador is mostly done from nests of the three genera selected here *Geotrigona* Moure, 1942, *Melipona*, Illiger, 1806 and *Scaptotrigona*, Moure 1943. The 20 pot-honey samples collected from cerumen pots and three ethnic honeys “abeja de tierra”, “bermejo”, and “cushillomishki” were analyzed for qualitative and quantitative ^1^H-NMR honey profiling and for the Honey Authenticity Test by Interphase Emulsion (HATIE). Extensive data of targeted organic compounds (41 parameters) was identified, quantified and described. The three honey types were compared by ANOVA. Amino acids, ethanol, hydroxymethylfurfural, aliphatic organic acids, sugars, and markers of botanical or entomological origin. The number of phases observed with the HATIE was one in *Scaptotrigona* and three in *Geotrigona* and *Melipona* honeys. Acetic acid (19.60 ± 1.45 g/kg) and lactic acid (24.30 ± 1.65 g/kg) were particularly high in *Geotrigona* honey (in contrast to 1.3 g/kg acetic acid and 1.6 g/kg lactic acid in *Melipona* and *Scaptotrigona*), with the lowest fructose + glucose (18.39 ± 1.68) g/100g honey compared to *Melipona* (52.87 ± 1.75) and *Scaptotrigona* (52.17 ± 0.60). Three local honeys were tested using PCA (Principal Component Analysis), two were assigned with a correct declared bee origin, but “bermejo” was not a *Melipona* and grouped with the *Scaptotrigona* cluster. However, after HCA (Hierarchical Cluster Analysis), the three kinds of honey were positioned in the *Melipona*-*Scaptotrigona* cluster. This research supports the targeted NMR-based profiling in pot-honey metabolomics approach for multi-parameter visualization of organic compounds, descriptive and pertained multivariate statistics (Hierarchical Cluster Analysis HCA, and Principal Component Analysis PCA) to discriminate the stingless bee genus in a set of *Geotrigona*, *Melipona* and *Scaptotrigona* honey types. The NMR characterization of Ecuadorian honey produced by stingless bees is a contribution to the needed regulatory norms. A final note on searching stingless bee markers in pot-honey metabolites that may become nutritional trait candidates for phylogeny. *Scaptotrigona* honey revealed biosurfactant activity in the HATIE, originating a fingerprint Honey Biosurfactant Test (HBT) for the genus in this set of pot-honeys.

## 1. Introduction

The tribe Meliponini (Hymenoptera, Apidae, Apinae) is the stingless bee entomological group (Michener, 2000) known to store and process honey and pollen in cerumen pots. The scientific literature refers to their honey as SBH (stingless bee honey). The term pot-honey was proposed to drive attention to the interactions of nectar to honey transformations within the cerumen container made up of stingless bee wax and plant resins (Vit et al., 2013). Their peculiar chemical composition with higher water content and free acidity than honey extracted from *Apis mellifera* honeycomb, analytical comparisons were initiated with *Melipona* and *Scaptotrigona* honeys extracted from cerumen pots in Brazil (Gonnet et al., 1964). After reviewing some 500 honeys from 66 species of stingless bees, Ávila et al. (2018) emphasize that meliponine honey positive health effects and market potential are an innovative beacon in food, pharmaceutical and cosmetic industries, in a world dominated by *A. mellifera* honey. Health promoting bioactive properties with bee-and-plant origin sustain the nutraceutical and medicinal uses of pot-honey (Pimentel et al., in press). According to the Codex Standard definition, honey is produced only by honey bees, and is described as “Honey consists essentially of different sugars predominantly glucose and fructose…” (CODEX STAN, 1987), and water. A precious but unregulated product (Braghini et al., 2021) with a first National norm for kelulut –all stingless bees in Malaysia (Department of Standards Malaysia, 2017) and the second from Argentina for ‘Yateí’ bees *Tetragonisca fiebrigi* (Secretaría de Regulación y Gestión Sanitaria y Secretaría de Alimentos y Bioeconomía, 2019).

Almost 400 species of stingless bees are Neotropical from the about 500 species distributed in the tropics and subtropics (Camargo and Pedro, 2007). Substantial discoveries were predicted by Michener (2013), and currently more than 600 stingless bee species thrive in: 1. The Neotropics (500) from 34.90°S in Montevideo, Uruguay up to 27.03°N in Álamos, Sonora, Mexico, 2. Africa (16) from 28.54°S in Eshowe, South Africa up to 18.00°N in Njala, Sierra Leone, and 3. Indo-Malaysia/Australasia (100) from 36.41°S in Australia up to 24.23 °N in Taiwan; exceptionally up to 4000 m.a.s.l. in Peru and Bolivia (Roubik and Vergara, 2021). The greatest global biodiversity of Meliponini is in Ecuador, 100 species in 8 km^2^ of the Amazonian forest located in the Yasuní Biosphere, (Roubik, 2018). Compared to *Melipona* (73 spp) and *Scaptotrigona* (22 spp), *Geotrigona* (22 spp) is less represented in biodiversity collections, see number of species in Camargo and Pedro (2013) chapter and *Melipona* review (Engel, 2021). Bees from the Cretaceous have a humanitarian role. Sustainable meliponiculture increases the resilience of small-scale stingless bee keepers, obtains good quality products, and sustains productive crops thanks to pollination (Baumung et al., 2021). Stingless bee management and honey processing by meliponicultors also imprints sanitary quality to this product (Heard, 2016).

The Mayan stingless bee keeping with *Melipona beecheii* is well documented in Mexico (Weaver and Weaver, 1981). Despite the enormous biodiversity of 200 species of stingless bees in Ecuador (Vit et al., 2018), their ancient records of meliponiculture are unknown. The honey produced by stingless bees has been widely relished in the tropics (Schwarz, 1948), and interested the Brazilian Court of the International Exhibition (1862) to display *Melipona* and *Trigona* bees, honey and wax (Smith, 1863), but the request of Meliponini presence in the Ecuadorian Pavillion of the Milan EXPO 1915 was declined. Honey pots have different volumes and shapes (Aguilar et al., 2013), generally *Melipona* has larger pots than *Scaptotrigona*, and *Geotrigona* has elongated sausage-like pots –using palynological descriptors for pollen morphology P/E (ratio polar/equatorial view of a pollen grain) a perprolate pot for the underground bee, subprolate for *Melipona* and spheroidal for *Scaptotrigona* (Vit P, personal observations), as illustrated in Figure 1.

**Figure 1.**
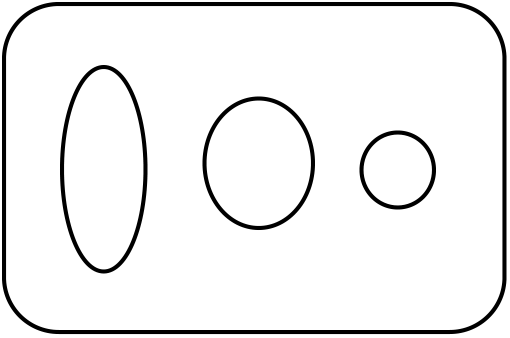
Diagram of equatorial view of honey pots from Ecuadorian *Geotrigona*, *Melipona* and *Scaptotrigona* from left to right.

Bacteria, molds and yeasts found naturally in honey are non-pathogenic, and new taxa of microbiota with commensal or mutualistic ecological role inside the stingless bee nests may offer sources of bioactive metabolites (Gilliam, 1997; Morais et al., 2013). Lactic acid is produced during the fermentation of honey (Vit et al., 2011). Little is known about relationships of honey and other products inside the stingless bee nest itself, e.g. *Tetragonisca angustula* from Venezuela (Pérez-Pérez et al., 2013), conversion of raw nectar into pot-honey (Leonhardt and Kaltenpoth, 2014). A seminal metabolomics approach investigated materials of the stingless bee nest: propolis, cerumen and honey produced by *Meliponula ferruginea* from Tanzania, storage modifications and comparison with *A.mellifera* (Popova et al., 2021), and supports future scientific research to answer how pot-honey is produced and kept within the cerumen pots.

Ecuadorian *A. mellifera* honey marketed in Quito (Salvador et al. 2019) and pot-honeys from twelve species of stingless bees (Villacrés-Granda et al., 2021) were assessed for physico-chemical quality. Nuclear magnetic resonance (NMR) based metabolomics was approached to fingerprint the entomological origin (Vit et al., 2015, Razali et al., 2018) and to detect adulteration (Yong et al., 2022) of meliponine honey. The advantage of NMR spectroscopy is the chemical profiling of a number of organic functional groups, to simultaneously quantify mono-, di-, trisaccharides, amino and organic acids, nucleobases, ethanol, HMF, and other characteristic constituents in a honey spectra (Popova et al., 2021). A systematic comparison between physico-chemical parameters and corresponding targeted NMR-based profile was done as a control for NMR in Chilean honeys (Fuentes Molina, et al., 2020), and Iranian honeys (Khansaritoreh et al., 2021). The large number of organic compounds identified and quantified by targeted NMR expands descriptive categories as a complement to the honey quality control analyses. Walker et al. (2022) consider the challenge of multiple analytical techniques leading to complex reports on sophisticated honey adulteration.

The aim of this study is to apply ^1^H NMR spectroscopy and a Honey Authenticity Test by Interphase Emulsion (HATIE) to compare the impact of the stingless bee genus in the organic matrix of their pot-honey. For this preliminary targeted metabolomics approach, twenty honey samples of *Geotrigona*, *Melipona* and *Scaptotrigona* were characterized with the content of 10 sugars, HMF and ethanol, 10 aliphatic organic acids, 10 amino acids, and 9 markers of botanical origin. Analysis of variance (ANOVA), hierarchical cluster analysis (HCA) and principal component analysis (PCA) based on NMR profiles were used to detect similarities and differences between the honey produced by *Geotrigona*, *Melipona*, and *Scaptotrigona* in Ecuador, and to position three test pot-honey samples at genus-level in the database. An idea on entomological markers and searching phylogenetic traits in honey is given at the end. A genus approach to study the current role and interactionts of stingless bees and their associated microbiota on sugary resources is to further in our understanding of environmental factors and putative phylogenetic relatedness of honey.

## 2. Materials and Methods

### 2.1 Pot-honey sampling

The harvest of pot-honeys was done in seven provinces of Ecuador: El Oro, Loja, Manabí, Napo, Orellana, Pastaza, and Santa Elena (Figure 1). The three biomes were represented: Coast (El Oro, Manabí, Santa Elena), Andes mountains known as Sierra (Loja), and Amazonian forest (Napo, Orellana, Pastaza). Twenty samples of mature honey from stingless bee colonies were extracted by suction with a plastic syringe connected with a rubber tube, after cutting a circular section with a syringe needle, from sealed cerumen pots –this means the nectar or the honeydew has been transformed into honey, similarly to the operculated *Apis mellifera* honeycombs. Two samples of *Melipona* were collected by Kichwa. The honey was transported at environmental temperature and kept frozen in the dark until analysis in the laboratory.

### 2.2 Entomological sampling and identification

Samples of stingless bees were collected from the nests in isopropyl alcohol, when possible, and sent dried for entomological identification to Dr. S.R.M. Pedro, Biology Department, Universidade de São Paulo, Ribeirão Preto, Brazil. They belong to three genera: *Geotrigona* Moure, 1943; *Melipona* Illiger, 1806, and *Scaptotrigona* Moure, 1942.

### 2.3 NMR

In nuclear magnetic resonance (NMR) spectroscopy, the chemical shift is the resonant frequency of a nucleus relative to a standard in a magnetic field. Often the position and number of chemical shifts are diagnostic of the structure of a molecule.

#### 2.3.1 NMR reference sample and honey sample preparation

All chemicals used in NMR were of analytical grade (>99% purity). The Hl 44518 - QuantRefA-NMR-Tube 5mm, 600µL (Reference Sample for quantification in food applications) was the NMR mixture analysis. This reference sample was used with Wilmad 507-PP-7 NMR-tubes in combination with Bruker’s Honey-Profiling. Deuterium oxide (D_2_O) was supplied by Sigma-Aldrich (Steinheim, Germany), 2,2,3,3-d(4)-3- (trimethylsilyl)propionic acid sodium salt (TSP) by Alfa Aesar (Karlsruhe, Germany), potassium dihydrogen orthophosphate (KH_2_PO_4_), sodium hydroxide (NaOH), and hydrogen chloride (HCl) by Merck (Darmstadt, Germany), sodium azide (NaN_3_) by Fluka (Steinheim, Germany), and deionized water was supplied by Th. Geyer (Renningen, Germany).The NMR profiles were obtained for functional groups of honey compounds: sugars, organic acids, amino acids, alkaloids, and alcohols. The pot-honey sample preparation method was adapted from BrukerBiospin (BrukerBiospin, Rheinstetten, Germany).The homogenized honey samples (5 g) were diluted in 17.5 mL NMR-buffer (15.7 g KH_2_PO_4_, 0.05 g NaN_3_ in 1L deionized water). For neutralization of organic acids the pH was adjusted at 3.1 with 1M HCl and 1M NaOH. Then, 900 µl of the homogenized honey solution were mixed with 100 µL standard solution (deuterium oxide D_2_O containing 0.1% TSP 2,2,3,3-d(4)-3-(trimethylsilyl) propionic acid sodium salt), centrifuged at 14000 rpm for 10 min, and 600 µL of supernatant were transferred into a 5 mm × 178 mm NMR-tube for direct measurement.

#### 2.3.1 Targeted quantitative ^1^H-NMR spectra acquisition and processing

All NMR spectra were acquired immediately after honey dilutions on a Bruker Ascend TM 400 MHz FoodScreener (BrukerBiospin, Rheinstetten, Germany) equipped with a 5 mm PA BBI 400SI H-BB-D-05 Z probe and using Bruker SampleXpress (BrukerBiospin, Rheinstetten, Germany) for automatic sample change. ^1^H-NMR-spectra were recorded at 300 K using the pulse program noesygppr1d (1D spectra with water presaturation at 4.8 ppm) and jresgpprqf (2D J-resolved spectra, displaying chemical shift and spin-spin coupling information). For 1D spectra, 32 scans and 4 dummy scans of 64 k points were acquired with a spectral width of 20.55 ppm, a receiver gain of 16 and an acquisition time of 3.99 s. The 2D spectra were performed using 4 scans and 16 dummy scans of 8 k (F2-axis) and 40k (F1-axis) points. The spectral widths were 16.7 ppm (F2) and 0.19 ppm (F1), receiver gain of 16 and an acquisition times were 0.6 s (F2) and 0.3 s (F1). The NOESY (Nuclear Overhauser Effect Spectroscopy) spectra were used for quantification and the JRES (J-resolved spectrum) were used to verify the compound identifications. All spectra were standardized, automatically phased, baseline-corrected, and calibrated using 2,2,3,3-D4-3- (trimethylsilyl)propionic acid sodium salt (TSP) as a reference at 0.0 ppm. The compounds were quantified by using the Honey-Profiling routine (release 1.0, BrukerBiospin, Rheinstetten, Germany) by automatic integration of the peak area calculated with an external standard (Spraul et al., 2009).

### 2.4 Honey Authenticity Test by Interphase Emulsion (HATIE)

This method was protected by two patents (Vit, 1995, 1999). It was created to detect fake honey (Vit, 1998). A disposable syringe was used to measure 0.5 mL honey and dilute it with 0.5 mL distilled water, 1 mL of diethyl ether was added and shaked by up and down wrist movements for 20 sec., let stand still for 1 min. and the number of phases was observed and noted. It was repeated with a 10 mL cylinder using 1 g honey, diluted with 1 g distilled water, diethyl ether was added to duplicate the volume of the dilution, and vortexed. It was let stand still for 1 min. and the number of phases was observed and noted. Two patterns with one or three phases were reported for genuine honey with this test, and illustrated in Figure 3 (Vit, 2022). The unique phase of *Scaptotrigona* honey was produced by an active biosurfactant of suspected microbial origin that needs to be demonstrated.

**Figure 2.**
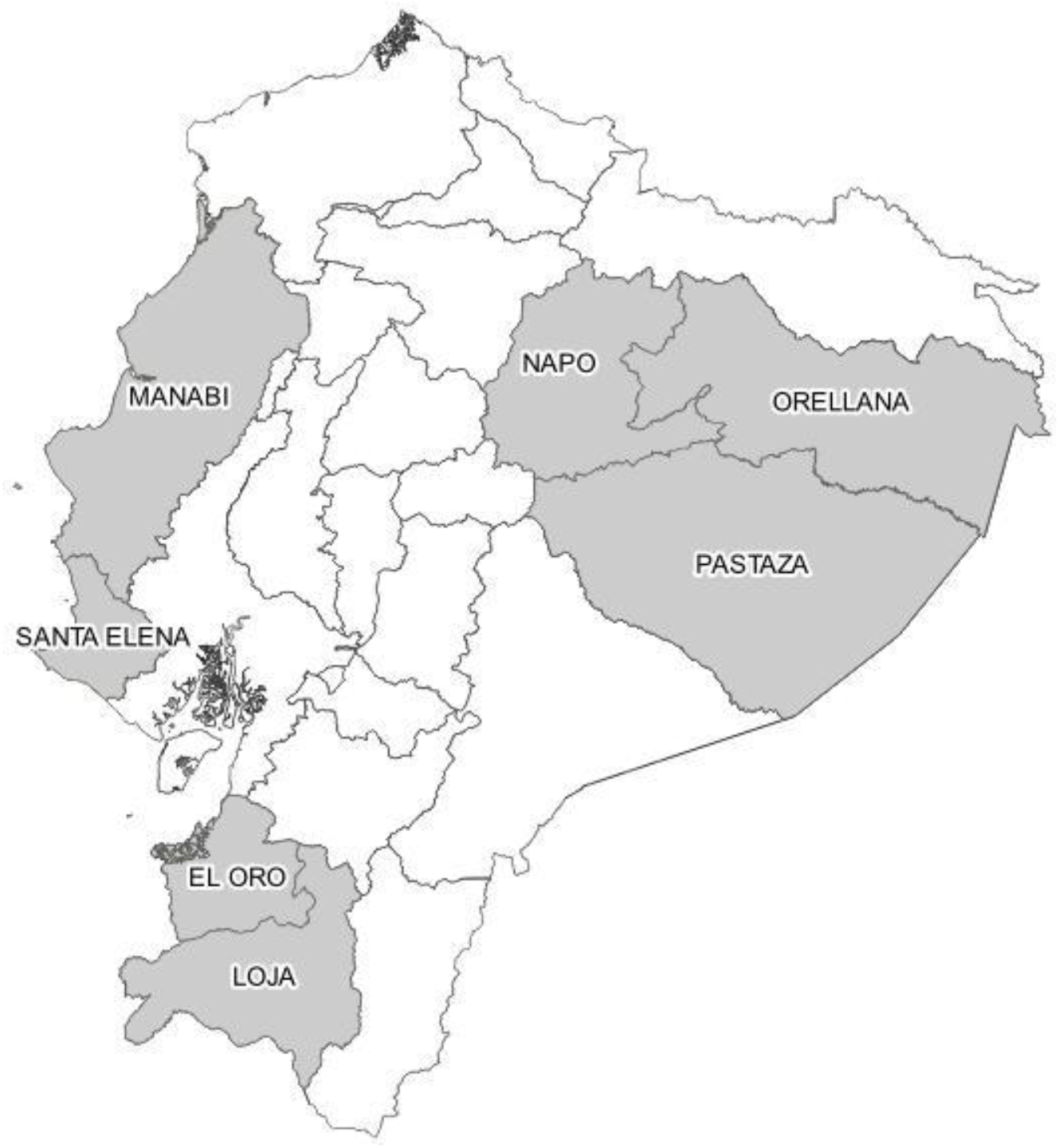
Locations of pot-honey harvest in seven provinces of Ecuador: **1.** El Oro, **2**. Loja, **3**. Manabí, **4**. Napo, **5**. Orellana, **6**. Pastaza, and **7**. Santa Elena

**Figure 3.**
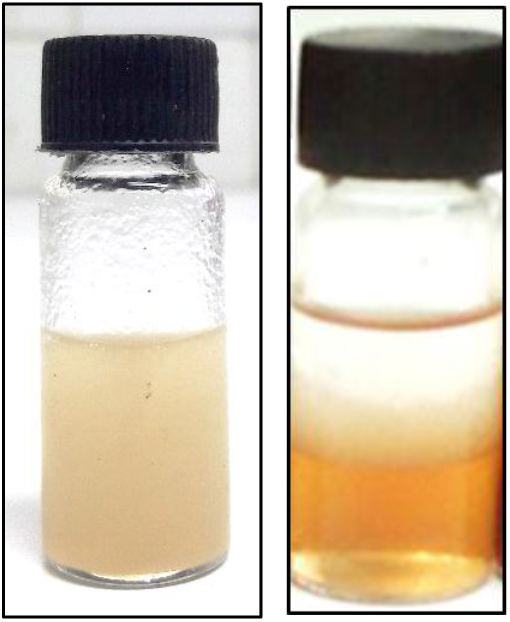
*Scaptotrigona*, fake and *Melipona* honey phases after the HATIE *From:* Vit (2022)

Vit (2022) presented a diagram for the number of phases in six Ecuadorian honey types: 1. One phase (*Scaptotrigona*), 2. Two phases (fake honey), and 3. Three phases (*Apis*, *Geotrigona*, *Melipona*, *Tetragonisca*).

### 2.5 Statistical analysis

The three entomological honey types were analyzed by ANOVA one way (P< 0.05), and post-hoc Scheffé test was perfomed to compare organic chemical components based on targeted NMR quantifications of pot-honey produced by *Geotrigona*, *Melipona* and *Scaptotrigona*, using the version 26 IBM SPSS Statistics software (IBM, 2019). Hierarchical cluster analysis (HCA) for sugars and organic acids were generated using the squared Euclidean distance between groups and using the Ward’s method. Pearson correlation matrix (P< 0.05), and Principal component analysis (PCA) were conducted using (XLSTAT Software Premium, 2021). Microsoft Power Point 2018 was used to prepare, integrate, and edit the figures composed from multiple sources.

## 3. Results and Discussion

The following 41 parameters (10 sugars, HMF and ethanol, 9 organic acids, 10 amino acids, and 10 markers of botanical or entomological origin) were quantified in the ^1^H NMR spectra of Ecuadorian pot-honey samples: 1. Sugars (fructose, glucose, sucrose, gentiobiose, maltose, maltotriose, mannose, turanose, mannose, melezitose, raffinose, and turanose), 2. HMF and ethanol, 3. Organic acids (acetic acid, citric acid, formic acid, fumaric acid, lactic acid, malic acid, pyruvic acid, quinic acid, and succinic acid), 4. Amino acids (alanine, aspartic acid, glutamine, isoleucine, leucine, phenylalanine, proline, pyroglutamic acid, tyrosine, valine), 5. Markers of botanical origin (acetoin, 2,3-butanediol, dihydroxyacetone, kynurenic acid, methylglyoxal, methylglyoxal dihydrate, methylglyoxal monohydrate, 3- phenyllactic acid, shikimic acid, and trigonelline). Quinic acid, isoleucine and kynurenic acid were not detected in the twenty honeys analyzed.

In the study of commercial Manuka honey the aliphatic components of honey (lactic acid, acetaldehyde, methylglyoxal dihydrate and methylglyoxal monohydrate, pyruvaldehyde, and pyruvic acid) showed signals in the first spectroscopic region (1.3-2.3 ppm), and phenyllactic acid in the aromatic region (6.0-8.0 ppm) of the ^1^H NMR spectra (Gresley et al., 2012). In a comparison of commercial Ecuadorian *Apis mellifera*, *Geotrigona* and *Scaptotrigona* honeys, a central region (3.5-4.5 ppm) of the ^13^C NMR spectra was characteristic for each bee genus, because the endogenous lipophilic markers of the bees obtained with chloroform extracts are independent of floral and geographical origin (Vit et al., 2015), as described.

The organic chemical profile (41 compounds) for the honey produced by three genera of stingless bees has remarkable differences and similarities. The variability of the honey they produce is a biodiversity trait. The quantitative data presented in the following Tables 1-5 will be described and discussed accordingly. Targetted ^1^H NMR is is a powerful technique for multiparametric analysis. Qualitative identification of selected metabolites in the reference standard and their corresponding quantitative measurements are possible. In the Supplementary Table 1S are given the NMR ranges (ppm) and type of signal for each metabolite.

**Table 1.** Sugar contents in Ecuadorian pot-honey quantified with ^1^H NMR.

| Sugars<br>(g/100g honey) | Stingless bee genus |  |  |
| --- | --- | --- | --- |
|  | *Geotrigona*<br>(n=8) | *Melipona*<br>(n=4) | *Scaptotrigona*<br>(n=8) |
| Fructose | $10.58 \pm 0.79^a$<br>[7.83 - 13.65] | $29.53 \pm 1.84^b$<br>[25.36 - 33.93] | $29.85 \pm 0.49^b$<br>[27.92 - 31.81] |
| Glucose | $7.81 \pm 0.94^a$<br>[4.9 - 11.92] | $23.23 \pm 0.79^b$<br>[21.65 - 25.30] | $22.33 \pm 0.29^b$<br>[21.31 - 23.42] |
| Sucrose | $0.15 \pm 0.02^a$<br>[0.07 - 0.28] | $0.10 \pm 0.05^a$<br>[0.02 - 0.24] | $0.20 \pm 0.06^a$<br>[0.01 - 0.45] |
| Gentiobiose | $0.20 \pm 0.07^a$ | $0.02 \pm 0.01^a$ | $0.17 \pm 0.11^a$ |
|  | [0.00 - 0.44] | [0.01 - 0.04] | [0.01 - 0.89] |
| Maltose | $0.49 \pm 0.05^a$<br>[0.27 - 0.69] | $0.83 \pm 0.32^a$<br>[0.27 - 1.47] | $2.02 \pm 0.21^b$<br>[0.92 - 2.75] |
| Maltotriose | $0.10 \pm 0.01^a$<br>[0.03 - 0.15] | $0.05 \pm 0.02^a$<br>[0.01 - 0.09] | $0.10 \pm 0.01^a$<br>[0.03 - 0.14] |
| Mannose | n.d. | $0.04 \pm 0.02$<br>[0.00 - 0.08] | n.d. |
| Melezitose | $0.07 \pm 0.00^a$<br>[0.05 - 0.08] | $0.12 \pm 0.03^a$<br>[0.08 - 0.21] | $0.25 \pm 0.03^b$<br>[0.16 - 0.38] |
| Raffinose | $5.03 \pm 0.31^b$<br>[4.13 - 6.67] | $0.45 \pm 0.06^a$<br>[0.34 - 0.62] | $1.24 \pm 0.09^a$<br>[0.88 - 1.69] |
| Turanose | $0.64 \pm 0.05^a$<br>[0.37 - 0.78] | $0.36 \pm 0.05^a$<br>[0.27 - 0.50] | $1.58 \pm 0.28^b$<br>[0.00 - 2.54] |
| **Total quantified sugars** | **25.07** | **54.73** | **57.75** |
| Fructose + Glucose | $18.39 \pm 1.68^a$<br>[13.08 - 24.50] | $52.87 \pm 1.75^b$<br>[49.12 - 56.10] | $52.17 \pm 0.60^b$<br>[49.71 - 54.96] |
| Fructose/Glucose | $1.42 \pm 0.10^a$<br>[1.05 - 2.06] | $1.27 \pm 0.11^a$<br>[1.07 - 1.57] | $1.34 \pm 0.03^a$<br>[1.24 - 1.48] |

Results are expressed as means ± standard error and [minimum - maximum] values. n.d. not detected. Different superscripts in a row show significant difference between G-M-S honeys with ANOVA (Scheffé, $P < 0.05$ ).

### 3.1 Sugars

The signals of glucose and fructose have a key role to discriminate *A. mellifera* honey (Lolly et al., 2008; Ohmenhaeuser et al., 2013). It is essential overcoming the honey matrix interference and the two main sugars with much higher peak intensities that may cover remaining signals of other compounds. The targeted sugar profile consisted of monosaccharides (fructose, glucose, and mannose), disaccharides (gentiobiose, maltose, sucrose, and turanose), and a trisaccharides (maltotriose, melezitose glucose, and raffinose galactose). The contents of the ten targeted sugars, the fructose + glucose value and the fructose/glucose ratio (Table 1) have differences and similarities between the three honey types. The total content of sugars (g/100 g honey) varied from 25.07 *Geotrigona* < 54.73 *Melipona* < 57.75 *Scaptotrigona*.

The ratio fructose/glucose (F/G) is used to predict slow crystallization for values < 1.3. This ratio was similar in the three genera (1.3-1.4), lower than the unprecedented above 2 reported for the Tanzanian *Meliponula ferruginea* (Popova et al., 2021), but similar to the Australian *Tetragonula carbonaria* (Persano Oddo et al., 2008). The F/G was 2.06 in one *Geotrigona* honey. However the low fructose content of *Geotrigona* (10.58 g/100 g) was almost triplicated in the other two genera, as well as the glucose content of *Geotrigona* (7.81 g/100 g) was triplicated in the *Melipona* and *Scaptotrigona* honeys, possibly indicating this underground bee has more active enzymes to catabolize fructose and glucose from the nectar, or its associated microbiota does it. Consequently the fructose + glucose was also lower in the *Geotrigona* (18.4) than *Melipona* (52.9) and *Scaptotrigona* (52.2). Sucrose was not statistically different but *Melipona* (0.10) < *Geotrigona* (0.15) < *Scaptotrigona* (0.20). Very low values compared to the maximum 5% permitted for *A. mellifera* honey (CODEX STAN, 1987). Average gentiobiose varied from 0.02 in *Melipona* and 0.17 to 0.20 in the other two honey types.

Persano Oddo et al. (2008) observed a putative maltose (15.3-22.8 g/100 g) of the Australian *Tetragonula carbonaria* honey not perfectly coincident with the maltose HPLC maltose standard, and suggested a further investigation, similar to the previous observations (2.5-3.2 g/100 g *Melipona* and 19.7-32.3 g/100 g other genera) by Bogdanov et al. (1996) in Venezuelan pot-honeys. Therefore, the maltose content of *Tetragonula carbonaria* reported in 2008, could correspond to the trehalulose recently discovered by Fletcher et al. (2020) in pot-honey harvested from four genera of stingless bees in Australia (*Tetragonula*), Brazil (*Tetragonisca*), and Malaysia (*Geniotrigona* and *Heterotrigona*). A new international regulation for honey including trehalulose is mandatory (Zawawi et al., 2022), and also the inclusion of trehalulose in the reference sample for targeted NMR Bruker’s Honey-Profiling. However the trehalulose content of 28.38 g/100 g honey for the Australian and Malaysian species, and the maltose content that varied from 0.49 to 2.02 g/100g in the Ecuadorian honey would not match with that quantity. It will not match either with the lower quantities of maltose in *Melipona* and *Scaptotrigona* species from Venezuela, it could similar to values of other stingless bees (*Frieseomelitta nigra paupera*, *Nannotrigona* aff. *chapadana, Tetragonisca angustula*) but it was higher for two *Nannotrigona* aff. *varia* (38.7-56.4) g/100 g honey, with corresponding fructose (16.1-13.9) and glucose (13.9-6.7) (Vit et al., 1998a). Maltose content of *Meliponula ferruginea* honey from Tanzania was low (1.56) and trehalulose lower than that (0.35), their fructose (26.9) high and lower glucose (12.8) causing the distinctive F/G >2 (Popova et al., 2021). The fructose + glucose content was intermediate to Ecuadorian honey, approximately 20 < 30 < 40 < 50 g/100g comparing *Geotrigona* < Australia-Malaysian *Geniotrigona*-*Heterotrigona-Tetragonula* < African *Meliponula* < Neotropical *Melipona* and *Scaptotrigona*. The trehalulose was also suggested as an additional marker of honeydew honey in Spain, instead of quercitol (de la Fuente et al., 2007), and similarly in some Bulgarian and Macedonian honeydews (Gerginova et al., 2020). Some statistical differences are significant for a higher maltose in *Scaptotrigona* (2.02), but the ranges of individual honeys overlap between *Melipona* [0.27 - 1.47] and *Scaptotrigona* [0.92 - 2.75].

The remaining five sugars were quantified in minor concentrations. Mannose was detected only in *Melipona* (0.04) and not in the other two genera. A lower maltotriose in *Melipona* (0.05), higher melezitose in *Scaptotrigona* (0.25), higher raffinose in *Geotrigona* (5.03), and higher turanose in *Scaptotrigona* (1.58). If the post-hoc ANOVA is done with Duncan test, less conservative than the Scheffé we used, raffinose is the unique sugar to differentiate the honey produced in our collection by *Geotrigona*, *Melipona* and *Scaptotrigona*. The total quantified sugars (g/100 g) varied from 25.07 for *Geotrigona*, 54.73 for *Melipona* and 57.75 for *Scaptotrigona*, a hallmark for pot-honey compared to higher concentrations of sugars in *A. mellifera* honey –a suggested standard of reducing sugars above 50g/100g for *Melipona* and *Scaptotrigona* is valid for the fructose + glucose values in Table 1 (Vit et al., 2004). The melezitose could be an indicator of honeydew honey (Persano-Oddo and Piro, 2004), and the raffinose too (Consonni et al., 2012). Interestingly, positive Pearson correlations were found between contents glucose and fructose (0.968), maltose and melezitose (0.853), maltose and turanose (0.801). Negative correlations were observed between raffinose and fructose (− 0.941), and glucose (−0.970). Raffinose (consists on galactose, glucose, and fructose) can be hydrolyzed to sucrose and galactose by the enzyme α-galactosidase (α-GAL). This may explain its negative Pearson correlations with fructose and glucose. Raffinose family of oligosaccharides (RFOs) are α-1, 6-galactosyl extensions of sucrose. This group of plant oligosaccharides is known to serve as desiccation protectant in seeds, as a transport sugar in phloem sap and as a storage sugar. Humans lack the α-GAL enzyme to break down this trisaccharide in their small intestine.

Nineteen sugars from aqueous extracts were identified in Italian honey of acacia, chestnut, rhododendron, multifloral and mountain by NMR: fructose, glucose, gentiobiose, isomaltose, kojibiose, maltose, maltulose, melibiose, nigerose, palatinose, sucrose, turanose, erlose, isomaltotriose, kestose, maltotriose, melezitose, raffinose, and maltotetraose (Consonni et al., 2012). Including also 11 unknown sugars, these authors suggested characteristic sets as identity cards for the Italian honeys: acacia (sucrose, turanose, fructose), chestnut (nigerose, kojibiose, raffinose, isomaltose, isomaltotriose, gentiobiose), rhododendron (melezitose, α-glucose, maltose, erlose, kestose), multifloral (β-glucose), and mountain (melezitose, α-glucose). Jang et al. (2016) studied 23 sugars (16 disaccharides and 8 trisaccharides) in Korean acacia, chestnut, multifloral, and artificial honey. Raffinose was more frequent and abundant in chestnut honey (0.01-0.17) mg/100 g.

The trisaccharide rafinnose quantified in honey by NMR (see Table 1) was used to create a dendrogram, where a separation of *Geotrigona* from a cluster of *Scaptotrigona* and *Melipona* honeys was achieved (See Figure 5B). *Melipona* and *Scaptotrigona* pot-honeys were separated by discriminant analysis of ten quality factors on Venezuelan species using only reducing sugars, sucrose, diastase activity and nitrogen, after factor reduction (Vit et al., 1998). Recently (Ávila et al., 2019) confirmed two distinctive clusters of Brazilian *Melipona* and *Scaptotrigona* honey by PCA of normalized mineral content and physicochemical analysis (moisture, Aw, electrical conductivity, total acidity, °BRIX, pH, colour, ash and total minerals). On a botanical approach, two clusters of *Heterotrigona itama* honeydew and blossom honey were revealed by PCA of physicochemical (ash content, hydrogen peroxide, free acidity, total mineral elements, K, Mg, Ca, total phenolics), and antioxidant (ferric reducing power) parameters in Malaysia (Ng et al., 2021). The honey produced by two species (*Tetragonula carbonaria* and *Tetragonula hockingsi*) of Australian stingless bees was separated from honey of two species (*Heterotrigona itama* and *Geniotrigona thoracica*) from Malaysia by HCA and PCA of moisture, electrical conductivity, pH, free acidity, color, glucose, fructose, trehalulose, total phenolics, total minerals, and 14 mineral elements (Zawawi et al., 2022).

**Figure 4.**
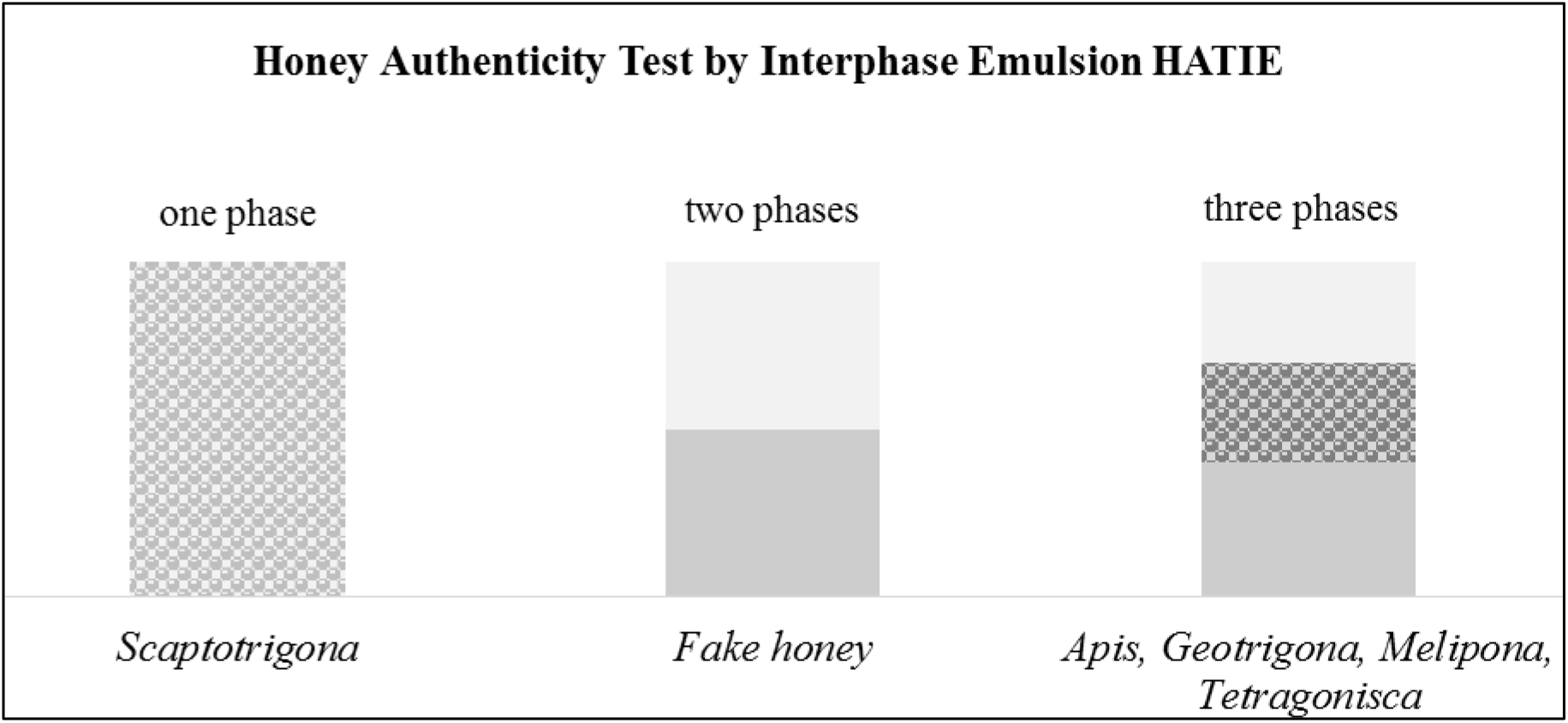
Number of phases in the honey test *From:* Vit (2022)

**Figure 5.**
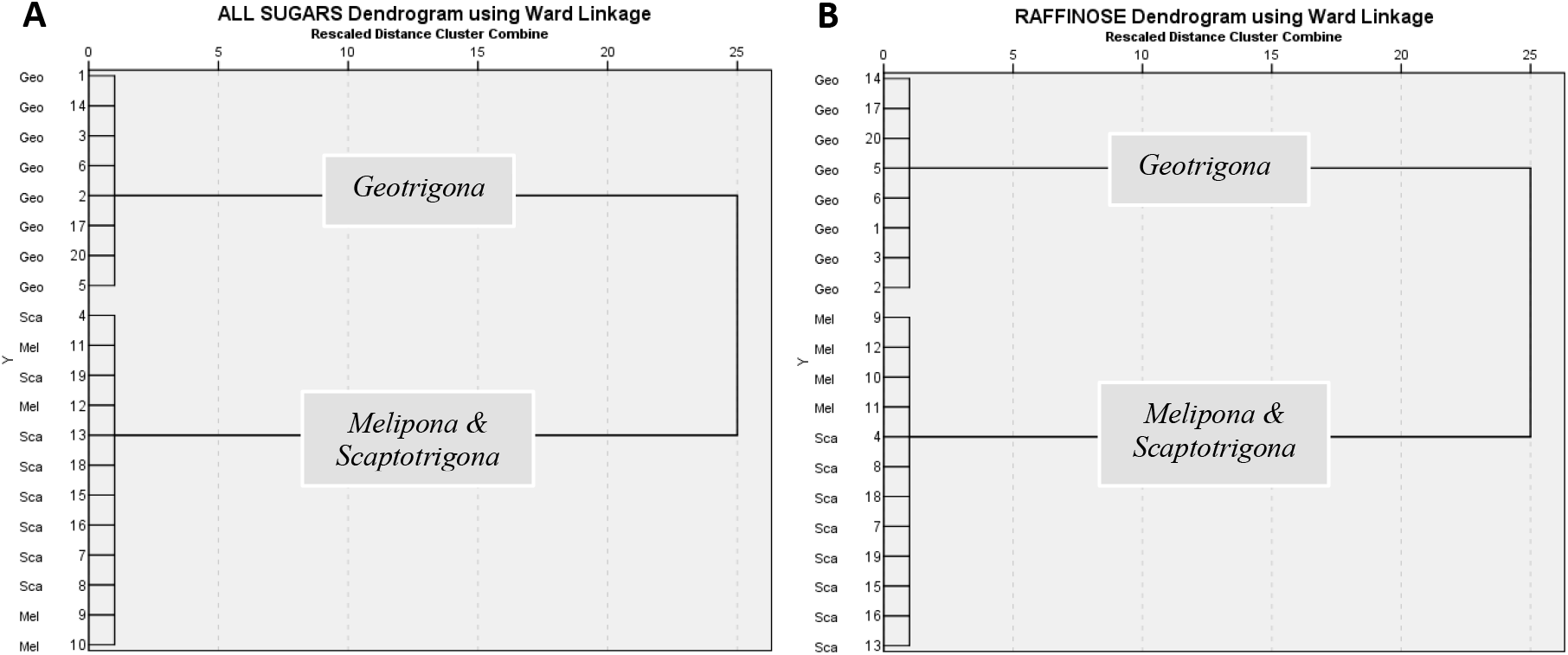
Dendrograms based on the content of total sugars **(A)** and raffinose **(B)** in honey produced by *Geotrigona* (Geo), *Melipona* (Mel) and *Scaptotrigona* (Sca). Observe that total sugars (A) generated alternated Mel-Sca honeys in the Mel & Sca cluster, but the raffinose (B) has ordered Mel first, followed by Sca in the root of the dendrogram. HCA by Euclidean distance using Ward’s method.

### 3.2 Hydroxymethylfurfural and ethanol

The hydroxymethylfurfural (HMF) is a degradation product of fructose caused by heating and aging, but also increases in acidic environments like that of honey. The presence of ethanol in honey derives from fermentation.

Both HMF and ethanol contents of honey are given in Table 2. The HMF content was higher in the *Geotrigona* (40.8 mg/kg) honey than the other two genera (7.2 *Scaptotrigona* and 20.9 *Melipona*), but ethanol was lower in this honey with substantial acetic and lactic acid concentrations caused by fermentation. The content of ethanol was also higher than *A. mellifera* in *Meliponula ferruginea* honey from Tanzania (Popova et al., 2021).

**Table 2.** HMF and ethanol contents in Ecuadorian pot-honey quantified with ^1^H NMR.

| Ethanol and HMF<br>(mg/kg honey) | Stingless bee genus |  |  |
| --- | --- | --- | --- |
|  | *Geotrigona* | *Melipona* | *Scaptotrigona* |

|  | (n=8) | (n=4) | (n=8) |
| --- | --- | --- | --- |
| **Ethanol** | 440.24 ± 100.60<sup>a</sup><br>[157.47 - 902.74] | 1553.66 ± 928.10<sup>a</sup><br>[17.56 - 3783.05] | 1753.17 ± 299.86<sup>a</sup><br>[998.5-3419.2] |
| **HMF** | 40.76 ± 5.81<sup>b</sup><br>[7.14 - 64.63] | 20.90 ± 13.70<sup>ab</sup><br>[0.00 - 60.21] | 7.21 ± 2.70<sup>a</sup><br>[1.64 - 19.71] |

Results are expressed as means ± standard error and [minimum - maximum] values. Different superscripts in a row show significant difference between G-M-S honeys with ANOVA (Scheffé, $P < 0.05$ ).

### 3.3 Aliphatic organic acids (AOA)

Natural aliphatic organic acids in honey are intermediates or final products of plant and microbiota Krebs cycle (citric, succinic, glutaric, fumaric, and oxaloacetic), fermentation (lactic, acetic), bee enzymes (gluconic), indicators of aerobic or anaerobic processes.

The content of AOA in pot-honey is given in Table 3. The total average content of nine organic acids varied from 0.29 to 4.43 g/100 g honey. Gluconic acid, lactic acid, and malic acid were studied in pot-honeys of *Tetragonula carbonaria* from Australia (Persano-Oddo et al., 2008) and *Melipona favosa* from Venezuela (Sancho et al., 2013), and from Malaysia (Shamsudin et al. 2019). Organic acids confer the organoleptic sour taste to honey and a distinctive acetic acid smell based on its concentration. Acetic acid and formic acid are volatiles identified in honey (Siegmund et al., 2017). All aliphatic organic acids were grouped in the targeted NMR approach of Chilean honeys (Fuentes Molina et al., 2020), here pyroglutamic acid is in the amino acids group, while kynurenic acid, 3-phenyllactic acid and shikimic acid are considered markers. Aliphatic organic acids in honey are of interest as indicators of fermentation or bioactivity. They were suggested as authenticity markers of bracatinga honeydew honey adulterated with floral honey from Brazil (Seraglio et al., 2021).

**Table 3.**
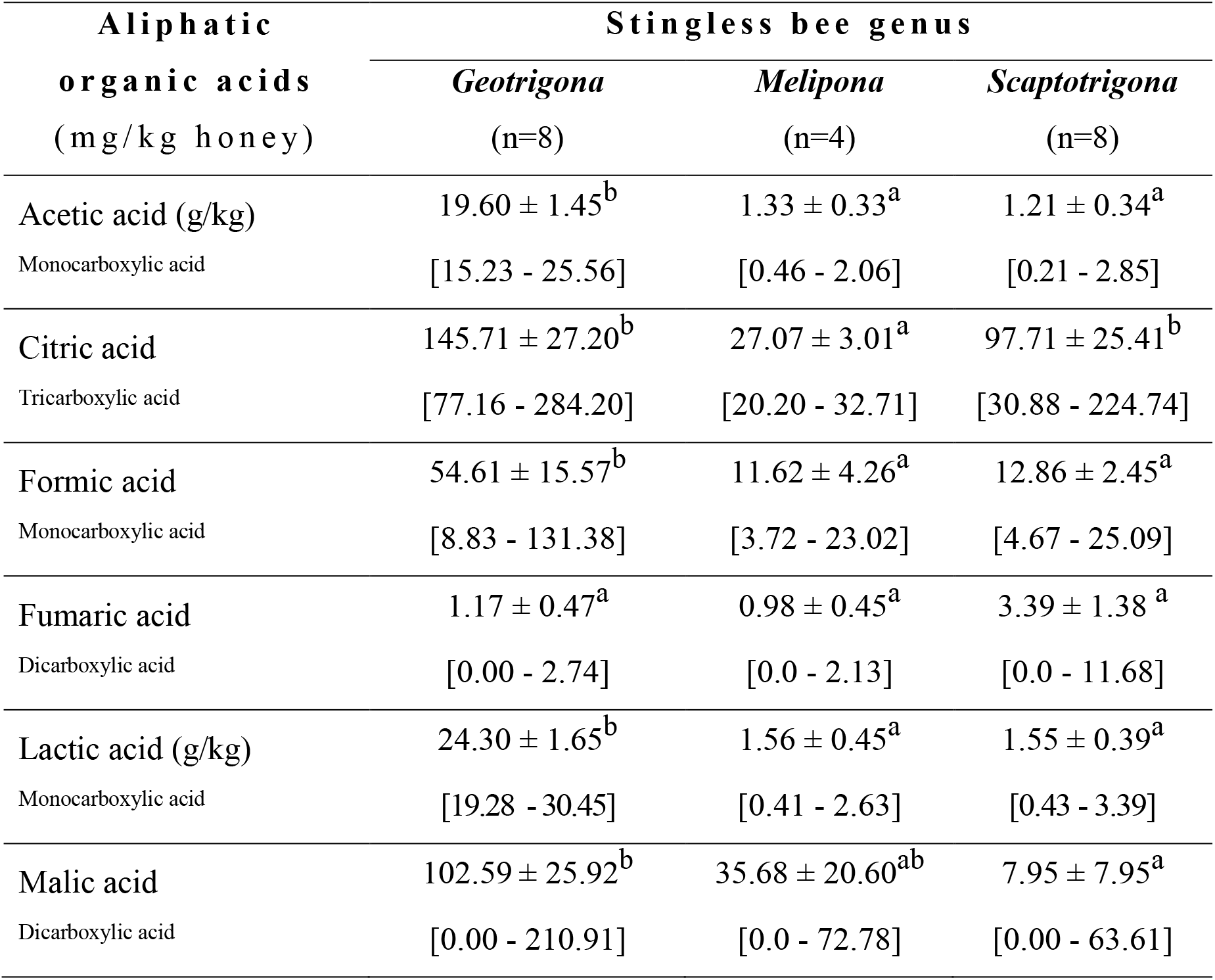
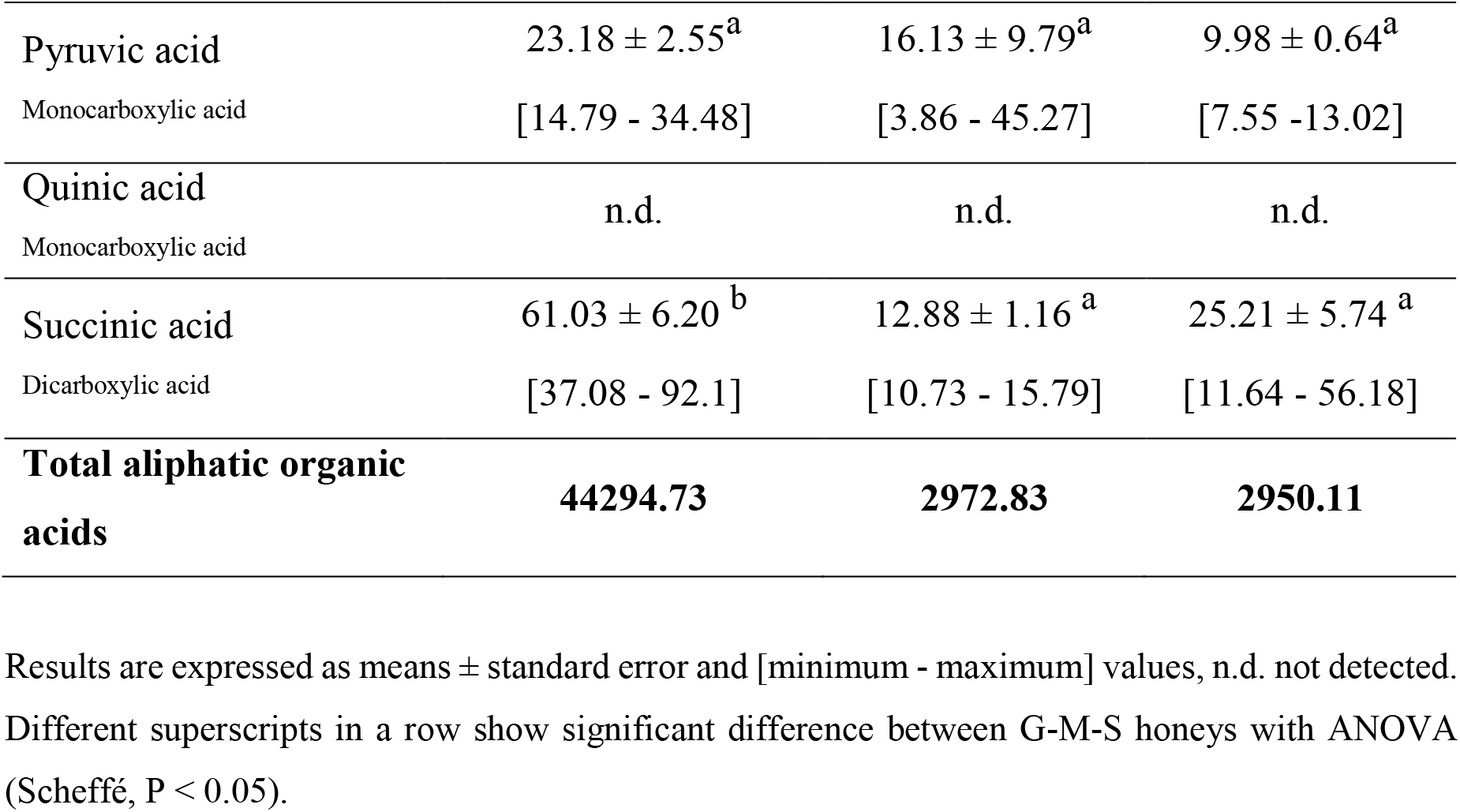
Aliphatic organic acid contents in Ecuadorian pot-honey quantified with ^1^H NMR.

| Aliphatic<br>organic acids<br>(mg/kg honey) | Stingless bee genus |  |  |
| --- | --- | --- | --- |
|  | *Geotrigona*<br>(n=8) | *Melipona*<br>(n=4) | *Scaptotrigona*<br>(n=8) |
| Acetic acid (g/kg)<br>Monocarboxylic acid | $19.60 \pm 1.45^b$<br>[15.23 - 25.56] | $1.33 \pm 0.33^a$<br>[0.46 - 2.06] | $1.21 \pm 0.34^a$<br>[0.21 - 2.85] |
| Citric acid<br>Tricarboxylic acid | $145.71 \pm 27.20^b$<br>[77.16 - 284.20] | $27.07 \pm 3.01^a$<br>[20.20 - 32.71] | $97.71 \pm 25.41^b$<br>[30.88 - 224.74] |
| Formic acid<br>Monocarboxylic acid | $54.61 \pm 15.57^b$<br>[8.83 - 131.38] | $11.62 \pm 4.26^a$<br>[3.72 - 23.02] | $12.86 \pm 2.45^a$<br>[4.67 - 25.09] |
| Fumaric acid<br>Dicarboxylic acid | $1.17 \pm 0.47^a$<br>[0.00 - 2.74] | $0.98 \pm 0.45^a$<br>[0.0 - 2.13] | $3.39 \pm 1.38^a$<br>[0.0 - 11.68] |
| Lactic acid (g/kg)<br>Monocarboxylic acid | $24.30 \pm 1.65^b$<br>[19.28 - 30.45] | $1.56 \pm 0.45^a$<br>[0.41 - 2.63] | $1.55 \pm 0.39^a$<br>[0.43 - 3.39] |
| Malic acid<br>Dicarboxylic acid | $102.59 \pm 25.92^b$<br>[0.00 - 210.91] | $35.68 \pm 20.60^{ab}$<br>[0.0 - 72.78] | $7.95 \pm 7.95^a$<br>[0.00 - 63.61] |
| Pyruvic acid | $23.18 \pm 2.55^a$ | $16.13 \pm 9.79^a$ | $9.98 \pm 0.64^a$ |
| Monocarboxylic acid | [14.79 - 34.48] | [3.86 - 45.27] | [7.55 -13.02] |
| Quinic acid | n.d. | n.d. | n.d. |
| Monocarboxylic acid |  |  |  |
| Succinic acid | $61.03 \pm 6.20^b$ | $12.88 \pm 1.16^a$ | $25.21 \pm 5.74^a$ |
| Dicarboxylic acid | [37.08 - 92.1] | [10.73 - 15.79] | [11.64 - 56.18] |
| **Total aliphatic organic acids** | **44294.73** | **2972.83** | **2950.11** |

Results are expressed as means ± standard error and [minimum - maximum] values, n.d. not detected. Different superscripts in a row show significant difference between G-M-S honeys with ANOVA (Scheffé, $P < 0.05$ ).

**Table 3a.**
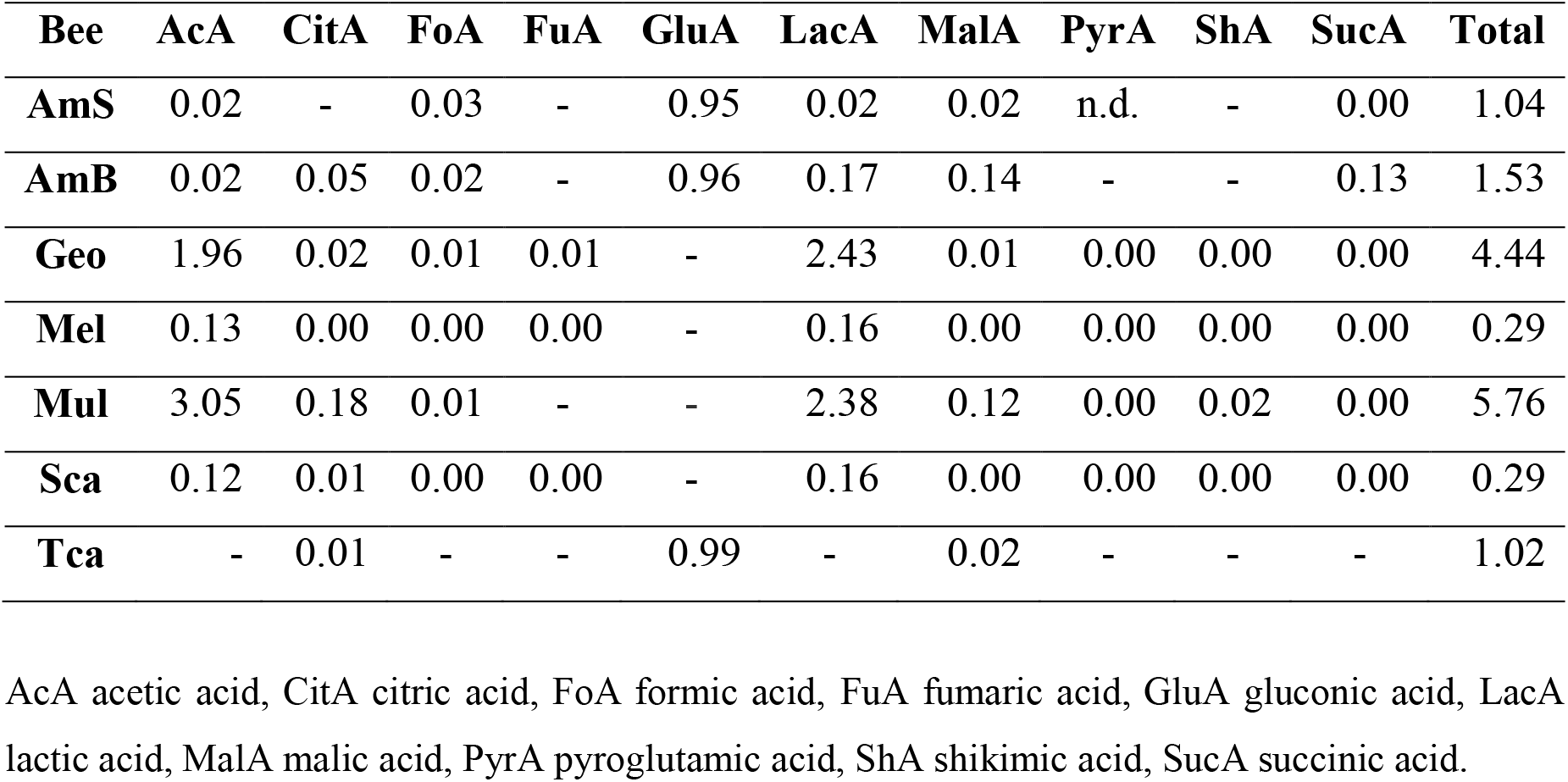
Contents of aliphatic organic acids (g/100 g) in *Apis mellifera* (AmS) honey from Spain (Mato et al., 2006), (AmB) from Brazil (Seraglio et al., 2021), and pot-honeys produced by *Tetragonula carbonaria* (Tcar) from Australia (Persano Oddo et al., 2008), *Meliponula ferruginea* (Mul) from Tanzania (Popova et al., 2021), *Geotrigona* (Geo), *Melipona* (Mel) and *Scaptotrigona* (Sca) from Ecuador in this work.

| Bee | AcA | CitA | FoA | FuA | GluA | LacA | MalA | PyrA | ShA | SucA | Total |
| --- | --- | --- | --- | --- | --- | --- | --- | --- | --- | --- | --- |
| AmS | 0.02 | - | 0.03 | - | 0.95 | 0.02 | 0.02 | n.d. | - | 0.00 | 1.04 |
| AmB | 0.02 | 0.05 | 0.02 | - | 0.96 | 0.17 | 0.14 | - | - | 0.13 | 1.53 |
| Geo | 1.96 | 0.02 | 0.01 | 0.01 | - | 2.43 | 0.01 | 0.00 | 0.00 | 0.00 | 4.44 |
| Mel | 0.13 | 0.00 | 0.00 | 0.00 | - | 0.16 | 0.00 | 0.00 | 0.00 | 0.00 | 0.29 |
| Mul | 3.05 | 0.18 | 0.01 | - | - | 2.38 | 0.12 | 0.00 | 0.02 | 0.00 | 5.76 |
| Sca | 0.12 | 0.01 | 0.00 | 0.00 | - | 0.16 | 0.00 | 0.00 | 0.00 | 0.00 | 0.29 |
| Tca | - | 0.01 | - | - | 0.99 | - | 0.02 | - | - | - | 1.02 |

AcA acetic acid, CitA citric acid, FoA formic acid, FuA fumaric acid, GluA gluconic acid, LacA lactic acid, MalA malic acid, PyrA pyroglutamic acid, ShA shikimic acid, SucA succinic acid.

In our study, acetic acid 19.60 g/kg and lactic acid 24.30 g/kg from *Geotrigona* were statistically higher than *Melipona* (1.33 and 1.56) and *Scaptotrigona* (1.21 and 1.55) honeys respectively. Indeed, the most remarkable sensory feature of *Geotrigona* honey is the very strong sour taste and acetic acid smell. Acetic (3.05 g/100 g) and lactic (2.38 g/100 g) acids for the Tanzanian *Meliponula ferruginea* honey (Popova et al., 2021), similar lactic acid but acetic acid above *Geotrigona* in our study. The family Acetobacteriaceae are acetic acid bacteria (AAB) and break down carbohydrates under acidic media, they occurred in *Tetragonula* but not in *Austroplebeia australis* (Leonhardt and Kaltenpoth, 2014a). Therefore, from our data, would be of interest to identify AAB in the Ecuadorian stingless bees.

Other organic acids are present at much lower concentrations in honey (mg/kg). Citric acid range was (20.6-284.2) and malic acid (0.0-210.9). The *Geotrigona* citric acid was almost triple of that in *Melipona* and double than *Scaptotrigona*. For malic acid *Geotrigona* (102.6) triplicated *Melipona* (35.7) in contrast with the low concentration found in *Scaptotrigona* (8.0). Three organic acids were evaluated in pot-honey of *Tetragonula carbonaria* from Australia by enzymatic methods: citric acid (0.23) and malic acid (0.12) were similar to *A. mellifera* honeys, but D-gluconic acid (9.9) was higher in *T. carbonaria* honey (Persano Oddo et al., 2008). In the honey of *Melipona favosa* from Venezuela citric acid (0.05) and lactic acid (0.03) (Sancho et al., 2013) were lower than the Ecuadorian *Melipona*. Gluconic acid is known to be the main acid responsible for the free acidity of *A. mellifera* honey and may indicate a high activity of glucose oxidase at ripening. Formic acid was also more concentrated in the *Geotrigona* (54.6) than *Melipona* (11.6) and *Scaptotrigona* (12.9), lower than values reported for *Apis mellifera* unifloral honeys 199-506 *Castanea*, 50-128 *Eucalyptus*, and 186-209 mg/kg multifloral (Suarez-Luque et al., 2006). *Geotrigona* and *Scaptotrigona* generate alarm signals near to their nests, but formic acid is used by *Oxytrigona* in caustic defensive secretion (Roubik, 1989). Formic acid, the simplest of fatty acids is the alarm pheromone in ant species (Blum and Brand, 1972). Although fumaric acid was higher in *Scaptotrigona* (3.4) than and *Geotrigona* (1.2) and *Melipona* (1.0), the variations caused no statistical difference between them. Pyruvic acid in *Geotrigona* (23.2) doubled the content of *Melipona* (16.1) and *Scaptotrigona* (10.0). Succinic acid was also higher in *Geotrigona* (61.0), some X5 of *Melipona* (12.0) and double of *Scaptotrigona* (25.2). Higher values were found in the honeydew honey from Brazil (0.5 to 0.7 g/100g (Brugnerotto et al., 2019). The quinic acid was absent in honey of the three genera, and the D-gluconic acid was not included in the NMR-based profiling, despite its importance in honey formation. It should be noted that gluconic acid has 16 stereoisomers, and has not been quantified in NMR-based profiles of honey so far because it is not easily measurable. D-gluconic acid can have theoretically 5 forms - one open, 2 pyranose, 2 furanose and all of them depending on pH could exist as acid or anion, all together 10 different forms in water (Simova S, personal communication). Gluconic acid accounted for 64.6 to 99.8% of total organic acids quantified by LC-MS/MS in *Apis mellifera* honey, and considered to be produced by worker bee enzymes such as glucose oxidase and the TCA cycle (Suto et al., 2020).

Lactic acid bacteria (LAB) microbiota are in symbiosis with *A. mellifera* (Olofsson et al. 2016) but Apini preserve their honey more dehydrated than Meliponini –fermentation by symbiont microbes such as *Zygosaccharomyces* and *Starmerella* species frequently isolated from their nests, have a role in pot-honey conservation (de Paula et al., 2021). Besides the LAB also acetic acid bacteria (AAB) are in symbiosis with the Australian *Tetragonula carbonaria* but not *Austroplebeia australis* (Leonhardt and Kaltenpoth, 2014). *Acetobacter*-like operational taxonomic unit (OTU) are associated with some stingless bees gut microbiota (Kwong et al., 2017). A cerumen pot made up of resins and stingless bee wax is rather soft and elastic, as needed to absorb the impact of anaerobic gas-related volume changes during nectar/honeydew fermentative processes to produce pot-honey (Vit P, personal observation), and additionally fibers of filamentous fungi evidenced in *Scaptotrigona depilis* brood cells (Menezes et al., 2015) may contribute to the material design with an organic holding mesh features strenght. A sealed honey pot is therefore, an ideal anaerobic container to maintain a wet environment in the warm nest temperature, needed for microbial proliferation in the sugary substrate harvested, carried and stored by the bees to fill open pots. *Bacillus cereus* was the most frequently isolated species in a *Melipona fasciata* nest from Panama (Gilliam et al., 1990), and *Bacillus* species were the major bacteria found in a *Heterotrigona itama* nest from Malaysia (Ngalimat et al., 2019). These spore forming bacteria secrete esterases, lipases, proteases, aminopeptidases, phosphatases, and glycosidases that transform food substrates into more digestible products. *Bacillus* species also secrete antibiotics and fatty acids to inhibit competing microorganisms which could spoil pot-honey or pot-pollen (Gilliam et al., 1990). The research of bee-microbes to protect pollinators may capitalize the discovery of novel bioactive compounds (Gilliam, 1997). Microbiome of stingless bees is highly variable. Wider bacteriological profiles of indoor surface are caused by a diversity of materials –of animal, plant, or soil origin– used by diverse species to build their nests (de Sousa, 2021).

A tentative set of aliphatic or non-aromatic organic acids is proposed for the *Geotrigona* exceptionally high free acidity and sour taste, in descending order of average concentration: 1. Lactic acid (24.3 g/kg), 2. Acetic acid (19.6 g/kg), 3. Citric acid (145.7 mg/kg), 4. Malic acid (102.6 mg/kg), 5. Succinic acid (61.0 mg/kg), and 6. Formic acid (54.6 mg/kg).

The organic acids quantified in honey (see Table 3) were used for HCA to create a dendrogram, where the *Geotrigona* cluster was separated from a cluster of *Scaptotrigona* and *Melipona* honey in Figure 6. Similar clusters were obtained in the dendrogram with all the sugars and the sugar raffinose, Figure 5.

**Figure 6.**
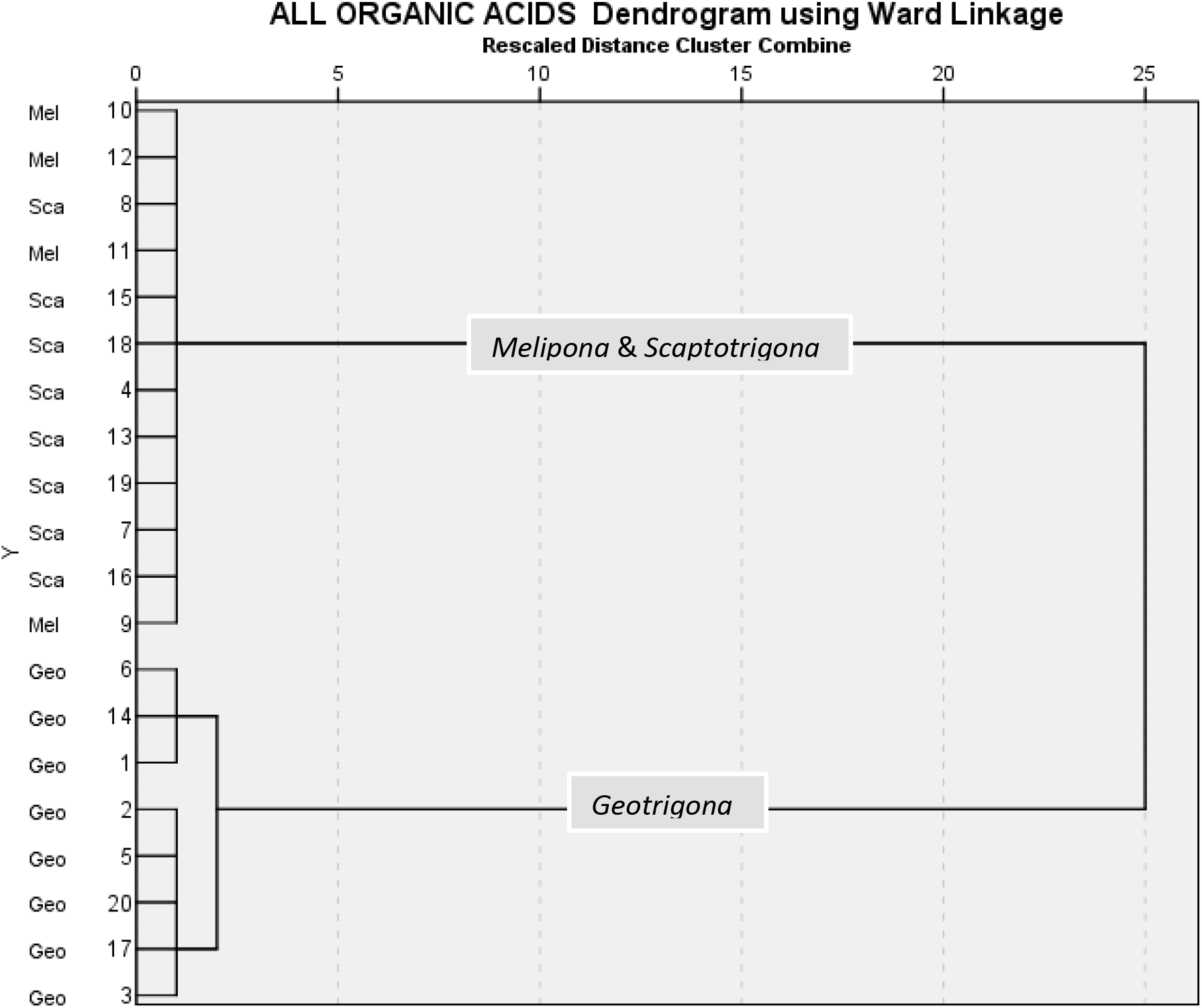
Dendrogram based on the content of all aliphatic organic acids in honey produced by *Geotrigona* (Geo), *Melipona* (Mel) and *Scaptotrigona* (Sca). HCA by Euclidean distance using Ward’s method.

In Table 3a harmonized units of g organic acid/100 g honey were used to express the data from Table 3 (g/kg and mg/kg) compared with the literature: 1. *Tetragonula carbonaria* (Tcar) from Australia (Persano Oddo et al., 2008) Unifloral and multifloral *Apis mellifera* (AmS) Spanish floral honey by capillary zone electrophoresis CZE (Mato et al., 2006), 2. *Apis mellifera* (AmB) Brazilian bracatinga honeydew honey by capillary electrophoresis CE (Seraglio et al., 2021) published as mg/100 g, 3. *Meliponula ferruginea* (Mul) honey from Tanzania, already using g/100 g (Popova et al., 2021). There are advantages and disadvantages in choosing any type of unit to express the content of organic acids in honey, because their range is wide. Using mg/kg is not practical for the hypervalues of most concentrated acids in pot-honey such as acetic acid and lactic acid. The visibility of g/100 g is reduced or even lost for the less concentrated acids, but these are usual units in food composition. Possibly mg/100 g honey is an intermediate option.

The total organic acids varied from 0.29 g/100 g honey for *Melipona* and *Scaptotrigona* in Ecuador to 5.76 for *Meliponula* in Tanzania (Popova et al., 2021), and 1.04-1.53 g/100g *Apis mellifera* honey in Table 3a. In the twelve species of stingless bees studied by (Villacres-Granda|, et al., 2021), the total organic acids measured by HPLC varied from 0.44 mg/100 g honey *Melipona mimetica* and 3.00 mg/100 g for *Tetragonisca angustula* from Ecuador. These authors measured acetic, citric, lactic and oxalic acids, Those values could be possibly in g/100 g honey). Their data is a great contribution with 27 pot-honeys, it was not only limited to all the honey quality standards, but also investigated the content of vitamin C, amino acids leucine and proline, organic acids, contaminant and nutritional metals, besides antibacterial activity. Acetic acid was the most abundant in *Scaptotrigona polysticta* and *Tetragonisca angustula* honey, but oxalic acid –a possible veterinary residue for *Varroa destructor* treatment of *Apis mellifera*, was the major organic acid in 10/12 stingless bee species. Its origin is unknown in pot-honey and may have a sensory impact on this already sour honey. In a study with eight Austrian unifloral honeys, a very unlikely set for floral volatiles (thujone isomers, eucalyptol, camphor, eugenol, thymol, and carvacrol) was suspected to originate in the use of essential oils by bee keepers (Siegmund et al., 2018). However, oxalic acid remained unchanged after varroa treatments (11 - 119 mg/kg) in Swiss honeys (Bogdanov et al., 2002), and similar values (14-114 mg/kg) in honeys from Spain (Mato et al., 2006).

From the eight organic acids studied by HPLC in *Geniotrigona thoracica* (Gt) and *Heterotrigna itama* (Hi) honey from Malaysia, formic acid was not detected (Shamsudin et al., 2019). These authors analyzed three monofloral types (acacia, starfruit and gelam) to observe further variations of organic acid contents (g/kg) caused by the botanical origin respectively: Acetic acid (0.09, 0.08, 0.06) Gt, (0.30, 0.06, 0.01) Hi; citric acid (0.04, 0.05, 0.03) Gt, (0.04, 0.06, 0.09) Hi; gluconic acid (0.48, n.d., 0.55) Gt, (0.90, 0.39, 1.48) Hi; lactic acid (0.20, 0.03, 0.17) Gt, (0.15, 0.18, 0.18) Hi; malic acid (0.30, n.d., 0.30) Hi only; succinic acid (0.52, 0.07, n.d.) Gt (0.32, 0.38, 0.34) Hi, and tartaric acid (0.04, n.d., n.d.) Gt, (0.06, 0.03, n.d.) Hi. The succinic acid was not present in Gelam honey, the other organic acids varied according to the bee species fed by different nectars. They found that gluconic acid was lower in starfruit than in the acacia and gelam honeys. Malaysian *H. itama* averages of floral and honeydew honey were similar and low for gluconic acid (0.47) and acetic acid (0.10) g/kg measured with enzymatic methods (Ng et al., 2021). The concentrations of these aliphatic organic acids given in (g/kg) should be multiplied by 10 to scale them into (g/100 g) as in Table 3a. All the Asian values will be very low compared with the African, Australian and South American pot-honeys, even for the gluconic acid that was measured enzymatically in Australian *T. carbonaria*, by CE in *A. mellifera* from Brazil and Spain, but not by NMR.

### 3.4 Amino acids

The origin of amino acids in honey is mostly undescribed in literature, except proline. The content of amino acids in pot-honey is given in Table 4. Proline and phenylalanine were the major amino acids in the three genera studied here, followed by alanine and pyroglutamic acid. In *Meliponula ferruginea* honey from Tanzania the major amino acid after proline was pyroglutamic acid, rarely found in *Apis mellifera* honey (Popova et al, 2021). In the Ecuadorian honeys proline varied from 34.41 to 565.77 mg/kg and averages were statistically different between *Geotrigona* (416.58) and *Scaptotrigona* (286.91), and *Melipona* (81.37) the lowest. Isoleucine was absent in all the honeys. The total average content of amino acids varied from 277.82 to 1183.10 mg/kg honey.

**Table 4.** Amino acid contents in Ecuadorian pot-honey quantified with ^1^H NMR.

| Amino acids<br>(mg/kg honey) | Stingless bee genus |  |  |
| --- | --- | --- | --- |
|  | *Geotrigona*<br>(n=8) | *Melipona*<br>(n=4) | *Scaptotrigona*<br>(n=8) |
| Alanine | $101.78 \pm 20.84^b$<br>[44.11 - 237.31] | $7.64 \pm 1.27^a$<br>[5.35 – 10.16] | $32.19 \pm 5.61^a$<br>[6.1-59.6] |
| Aspartic acid | n.d. | n.d. | $19.15 \pm 19.15$<br>[0.0-153.2] |
| Glutamine | 43.59 ± 11.22<sup>a</sup><br>[15.41 - 118.34] | 8.40 ± 0.99<sup>a</sup><br>[6.02 - 10.78] | 35.36 ± 14.36<sup>a</sup><br>[10.45 - 119.97] |
| Isoleucine | n.d. | n.d. | n.d. |
| Leucine | 22.57 ± 12.31<sup>a</sup><br>[0.0 - 93.73] | n.d. | 27.66 ± 8.79<sup>a</sup><br>[0.0-65.01] |
| Phenylalanine | 415.86 ± 82.37<sup>a</sup><br>[163.31 - 783.38] | 125.37 ± 56.90<sup>a</sup><br>[26.04 - 244.20] | 312.53 ± 79.80<sup>a</sup><br>[147.85 - 826.16] |
| Proline | 416.58 ± 43.83<sup>b</sup><br>[225.24 - 565.77] | 81.37 ± 28.77<sup>a</sup><br>[34.41 - 165.24] | 286.91 ± 39.05<sup>b</sup><br>[179.71 - 529.16] |
| Pyroglutamic acid | 96.03 ± 31.61<sup>a</sup><br>[0.00 - 218.26] | n.d. | 73.90 ± 49.05<sup>a</sup><br>[0.00 - 338.20] |
| Tyrosine | 56.11 ± 11.99<sup>a</sup><br>[25.62 - 124.62] | n.d. | 99.81 ± 24.28<sup>a</sup><br>[24.69 - 203.04] |
| Valine | 30.58 ± 5.20<sup>b</sup><br>[15.95 - 61.07] | 0.31 ± 0.31<sup>a</sup><br>[0.0 - 1.22] | 15.67 ± 3.91<sup>ab</sup><br>[3.72 - 37.17] |
| **Total average amino acids** | **1183.10** | **277.82** | **903.18** |

Results are expressed as means ± standard error and [minimum - maximum] values, n.d. not detected. Different superscripts in a row show significant difference between G-M-S honeys with ANOVA (Scheffé, $P < 0.05$ ).

Bumble bees do not differentiate between different amino acids, but between different concentrations of the same amino acid (Leonhardt et al., 2020). Indeed, amino acids are not a botanical marker candidate in unifloral and multifloral honeys (Hermosin et al., 2003; Carrafu et al., 2011), they were not either entomological markers for the three meliponine genera studied here, but L-alanine was suggested as an entomological marker for *Tetrigona apicalis* from Malaysia (Razali et al., 2018). Besides the bee, the nectar and the pollen sources, the amino acid spectra of pot-honey can vary by the action of symbiotic microbiota in the stingless bee nest (Barbosa et al. 2017). The *Monoascus* sp. fungal mycelia growing on the food supply within the brood cells is essential for the larval development of *Scaptotrigona postica* (Menezes et al., 2015).

Residual proline of honey originates from salivary secretions in cephalic glands of stingless bee workers added to transform nectar into honey. Proline content in honey of *Tetragonisca angustula* from Colombia was 770 mg/kg (Torres et al., 2004). Honeys of *Melipona scutellaris, Melipona quadrifasciata*, *Melipona subnitida,* and *Plebeia* sp. from Brazil varied between 201.6 and 924.6 mg/kg (Duarte et al. 2012). Similarly, honey of *Melipona beecheii* from Mexico varied from 264.5 to 1193.7 mg/kg (Moo-Huchin et al., 2015). Lower values were found in honey of *Melipona subnitida* and *Melipona scutellaris* from Brazil, ranging from 46.0 to 205.0 mg/kg (de Sousa et al., 2016). Biluca et al. (2019) analyzed 16 free amino acids (aspartic acid, glutamic acid, asparagine, glutamine, serine, arginine, glycine, threonine, alanine, proline, tyrosine, valine, leucine, isoleucine, phenylalanine, tryptophan) in honey of nine stingless bee species from Brazil, the two most concentrated and ubiquitous were phenylalanine (5.2-1231.0 mg/kg) and proline (12.1-762.0 mg/kg), glutamine (1-605) was absent in half of the honeys, histidine was absent in all. Valine (16.1-85.9) was present in all honeys except *Melipona bicolor*, like tyrosine (8.5-156.0) which was not present also in one of the two *Melipona quadrifasciata* honeys, possibly indicating a non entomological origin. The quantification of 17 amino acids (arginine, histidine, lysine, phenylalanine, isoleucine, leucine, methionine, valine, threonine, tyrosine, proline, glutamic acid, aspartic acid, serine, alanine, glycine, and cysteine) was done for the honey produced by *Tetragonula laeviceps* in three locations of Indonesia (Agussalim and Nurliyani, 2021). The average proline varied from 80.54 to 597 mg/kg, similar to the Ecuadorian pot-honeys ranges in Table 4 (34.4-565.8). Their higher concentrations were found for glutamic acid with 1238.74 (Sleman), 1922.98 (Lombok), and surprisingly it was ten fold lower in the third region (Gunungkidul), possibly related to the protein sources available. In the pot-honeys produced by twelve species of Ecuadorian stingless bees, proline concentration was higher than leucine, the other amino acid measured by Villacres-Granda|, et al., (2021). In their five *Melipona* spp. the average proline varied from 198.58 to 5132.71 mg/kg honey, in the *Scaptotrigona polysticta* it was 4179 mg/kg, and for the other species 84.56 -1459.51 mg/kg. In this study comparing three genera of Ecuadorian stingless bees, the highest average was 416.58 mg/kg for *Geotrigona*.

Pyroglutamic acid is a less known and studied uniquitous natural amino acid derivative in which the free amino group of glutamic acid or glutamine cyclizes to form a lactam. It was absent in *Melipona* honey and varied from 0.0 - 338.2 mg/kg with averages of *Geotrigona* (96.03) > *Scaptotrigona* (73.9).

### 3.5 Markers of plant and bee origin

Volatile markers of monofloral honey from different regions may vary according to the flora. Putative markers were the volatile compounds most often reported in twenty selected monofloral honeys produced by *A. mellifera* (Machado et al., 2020).

In Table 5, the nine targeted botanical markers quantified by ^1^H NMR (mg/kg), with the most concentrated 2,3-butanediol and the absent kynurenic acid. The shikimic acid is included in this table as a proposed entomological marker (Popova et al., 2021).

**Table 5.** Plant-bee markers in Ecuadorian pot-honey quantified with ^1^H NMR.

| Markers of botanical or<br>entomological origin<br>(mg/kg honey) | Stingless bee genus |  |  |
| --- | --- | --- | --- |
|  | *Geotrigona*<br>(n=8) | *Melipona*<br>(n=4) | *Scaptotrigona*<br>(n=8) |
| Acetoin | n.d. | $17.10 \pm 16.11^a$<br>[0.00 - 65.38] | $2.06 \pm 2.06^a$<br>[0.0-16.45] |
| 2,3-Butanediol | $502.31 \pm 112.13^a$<br>[101.32 - 972.82] | $264.29 \pm 141.78^a$<br>[15.47 - 530.16] | $127.59 \pm 37.82^a$<br>[28.79 - 323.83] |
| Dihydroxyacetone | $133.68 \pm 13.41^b$ | $32.06 \pm 7.39^a$ | $43.37 \pm 7.63^a$ |

|  | [90.89 - 183.83] | [23.01 - 54.15] | [14.76 - 85.64] |
| --- | --- | --- | --- |
| Kynurenic acid | n.d. | n.d. | n.d. |
| Methylglyoxal | n.d. | 3.27 ± 3.27<sup>a</sup><br>[0.0 - 13.18] | 4.38 ± 3.01<sup>a</sup><br>[0.0 - 22.37] |
| Methylglyoxal dihydrate | 18.45 ± 9.14<sup>a</sup><br>[0.0 - 58.15] | 19.66 ± 5.38<sup>a</sup><br>[7.85 - 43.51] | 9.47 ± 2.19<sup>a</sup><br>[0.00 - 20.64] |
| Methylglyoxal monohydrate | 21.96 ± 2.94<sup>b</sup><br>[14.14 - 39.29] | 6.54 ± 1.12<sup>a</sup><br>[3.56 - 8.59] | 5.74 ± 1.42<sup>a</sup><br>[1.92 - 12.76] |
| 3-Phenyllactic acid | 69.24 ± 34.80<br>[0.00 - 232.34] | n.d. | n.d. |
| Shikimic acid | 11.52 ± 3.43<sup>a</sup><br>[3.22 - 32.51] | 10.47 ± 3.85<sup>a</sup><br>[3.36 - 20.35] | 30.46 ± 14.45<sup>a</sup><br>[6.09 - 84.30] |
| Trigonelline | 18.94 ± 3.61<sup>a</sup><br>[5.64 - 32.47] | 2.16 ± 1.55<sup>a</sup><br>[0.00 - 6.59] | 25.32 ± 8.84<sup>a</sup><br>[6.09 - 84.30] |

Results are expressed as means ± standard error and [minimum - maximum] values, n.d. not detected. Different superscripts in a row show significant difference between G-M-S honeys with ANOVA (Scheffé, $P < 0.05$ ).

#### 3.5.1 Acetoin

Acetoin is 3-hydroxy-2-butanone, a natural volatile of the Australian *Eucalyptus leucoxylon* and *Eucalyptus melliodora* (D’Arcy et al., 1997). It is also present in heather honey from France (Radovic, 2000), Germany and Norway (Guyot et al., 1999). It was absent in *Geotrigona*. Only low concentrations were detected in *Melipona* (17.1) mg/kg and *Scaptotrigona* (2.1).

#### 3.5.2 2,3-Butanediol

The most concentrated botanical marker in the three Ecuadorian honeys was 2,3-butanediol, from 127.6 to 502.3 mg/kg honey being *Geotrigona* > *Melipona* > *Scaptotrigona*. This volatile wass found in rosemary and thymus honeys from Spain (Pérez et al., 2002), and was reported (mg/kg) in a sensory study of the Austrian unifloral honey volatiloma for dandelion 0.02, robinia 0.50, rape 0.40, fir tree 0.05, chestnut 0.17, linden 0.11, orange 7.77, and lavender 0.03 (Siegmund et al., 2018), all less concentrated than the Ecuadorian honey.

#### 3.5.3 Dihydroxyacetone

Dihydroxyacetone (DHA) is the precursor to methylglyoxal (Smallfield et al., 2018). Manuka phytochemicals including DHA were related with elements of the soil (Meister et al., 2021) Nectar chemical traits were different between sites, but plant-to-plant variation within sites were the largest (Icon et al., 2021). DHA varied from 32.1 to 133.7 mg/kg in the Ecuadorian pot-honeys, another botanical source is producing it because manuka is not growing in Ecuador. Glycerol is transformed to dihydroxyacetone by the bacterium *Acetobacter suboxidans* (Charney, 1978).

#### 3.5.4 Kynurenic acid

Resonances of minor compounds also play a certain role such as quinoline alkaloids and kynurenic acid for chestnut honey (Truchado et al., 2009). Kynurenic acid was found in small concentrations in Alpine honey from Italy (Leoni et al., 2021). In Moroccan jujube *Ziziphus lotus* honey kynurenic acid contents were 17.7 mg/kg by HPLC-DAD-ESI-MS and 6.6 mg/kg by GC-EI-MS (Khallouki et al., 2021). In the Ecuadorian pot-honeys, kynurenic acid was non-detected as in the Chilean honeys (Fuentes Molina et al., 2020).

#### 3.5.5 Methylglyoxal, Methylglyoxal dehydrate and Methylglyoxal monohydrate

These are markers of manuka (*Leptospermum scoparium*) honey. Antibacterial activity of honey after hydrogen peroxide release is known as non-peroxide activity (NPA) in Australia, similar to the Unique Manuka Factor (UMF) in New Zealand, with a rating >10+ considered therapeutically active (Cokcetin et al., 2016). These authors explain DHA is converted to MGO during storage, indicating the potential of honey to increase bioactivity. Highest MGO >800 mg/kg corresponded to highest NPA (21.9–26.3%). Methylglyoxal (MGO) was not detected in *Geotrigona* honey but in *Melipona* and *Scaptotrigona* it was 3.3-4.4 mg/kg respectively. However, both MGO dehydrate (9.5-19.7) and MGO monohydrate (5.7-22.0) were more concentrated in *Geotrigona*, but no significant differences between the three honey types.

#### 3.5.6 3-Phenyllactic acid

In thyme honey from Greece it varied from 1.5 to 28.1 g/kg (Alissandrakis et al., 2009). *Tetragonula carbonaria* raw honeys were harvested from “sugarbag” in Australia. The phenolic extracts averaged to 58.7 mg/kg and included 3-phenyllactic acid in a range from 3.7 to 12.7 mg of gallic acid equivalents/100 g honey (Massaro et al., 2014). Manuka (*Leptospermum scoparium*) and kanuka (*Kunzea ericoides*) nectars and honeys from New Zealand contain 3-phenyllactic acid, 740 mg/kg manuka honey and 660 mg/kg kanuka honey (Bong et al., 2018), which is a broad spectrum antimicrobial compound. In Table 5, 3-phenyllactic acid was found only in *Geotrigona* honey, 69.2 mg /kg.

#### 3.5.7 Shikimic acid

In microorganisms, shikimic acid is derived from phenolics, and the shikimate biosynthesis pathway produces precursors for the aromatic amino acids phenylalanine, tyrosine, and tryptophan. The physiology of the bee or the associated microbiota in the nest, also the bee diet, or a combination of them could explain the presence of shikimic acid in honey (Santos-Sánchez et al., 2019). A rare apple honey was characterized by high contents of shikimic acid-pathway derivatives (Kuś et al., 2013). In our study there is a wide variation from 3.22 - 84.30 mg/kg, as observed in Table 5, with a higher average in *Scaptotrigona* (30.5) than *Geotrigona* (11.5) and *Melipona* (10.5). Therefore it becomes interesting to describe the concentration of shikimic acid by *Scaptotrigona* in the honey they process, compared to that of *Geotrigona* and *Melipona*. However, variations were too high to separate these three honey types.

#### 3.5.8 Trigonelline

This nucleobase has been identified as coffee origin. In a study of unifloral honey from Colombia and multifloral from Honduras, trigonelline ranged from 23 to 86 mg/kg honey, and was suggested as a marker of *Apis mellifera* unifloral honey of *Coffea arabica* (Schievano et al., 2015). It was present in the *Meliponula ferruginea* honey from Tanzania (Popova et al, 2021), and averages here ranged from 2.2 to 25.3 mg/kg *Melipona* < *Geotrigona* < *Scaptotrigona*.

In the Chilean honey study (Fuentes Molina, et al., 2020), a reduced set of markers excluding the organic acids was named low molecular weight molecules, and they found it a quite heterogeneous group, like here. These markers did not separate the three genera of Ecuadorian honey.

In Figure 7, a graphic comparison is presented for selected targeted NMR based concentrations (from Tables 1, 3-5) of sugars (fructose, glucose, raffinose), organic acids (acetic acid, lactic acid, citric acid X10), amino acids (proline, phenylalanine, tyrosine) and plant markers (2,3-butanediol, dihydroxyacetone, trigonelline) in Ecuadorian honey produced by *Geotrigona*, *Melipona* and *Scaptotrigona*. Statistical differences by ANOVA Scheffé post-hoc were detected in parameters from the four chemical groups, sugars, organic acids, amino acids and markers. Therefore it is worth to continue with them, other sugars and alcohols would be needed in the reference standard if possible for Bruker.

**Figure 7.**
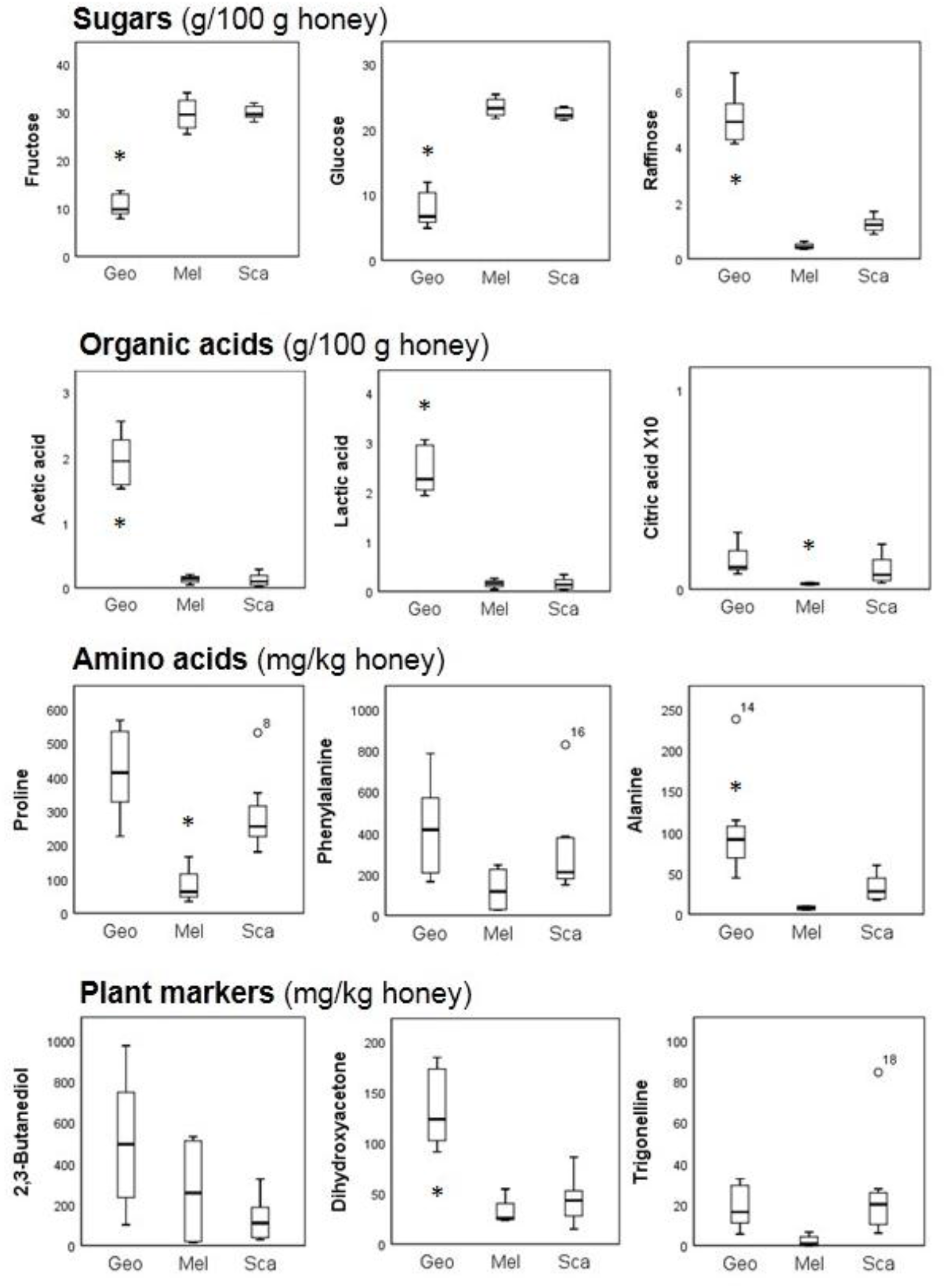
Major sugars (g/100 g), aliphatic organic acids (g/100 g), amino acids (mg/kg) and plant markers (mg/kg) in honey produced by *Geotrigona* (Geo) n=8, *Melipona* (Mel) n=4 and *Scaptotrigona* (Sca) n=8. Boxplots show the range of values (median thick line, lower 0.25 and upper 0.75 quartile white boxes, minimum and maximum values whiskers, and outliers of each dataset). Significant differences of honey between the three genera in each parameter were marked (Scheffé, * P< 0.05). Note that outliers in the three amino acids and trigonelline belong to different honey samples, one *Geotrigona* and three *Scaptotrigona*.

### 3.6 PCA Multivariate analysis

In Figure 8, the loading plots of active variables or honey components used in the PCA are organized for: 1. Sugars (8A), 2. Organic acids (8B), 3. Amino acids (8C), and 4. Markers of plant or bee origin (8D). In these plots F1 x F2, the variability of honey was explained 67.87% by sugars, 67.52% by organic acids, 59.78% by amino acids, and 48.84% by plant-bee markers.

**Figure 8.**
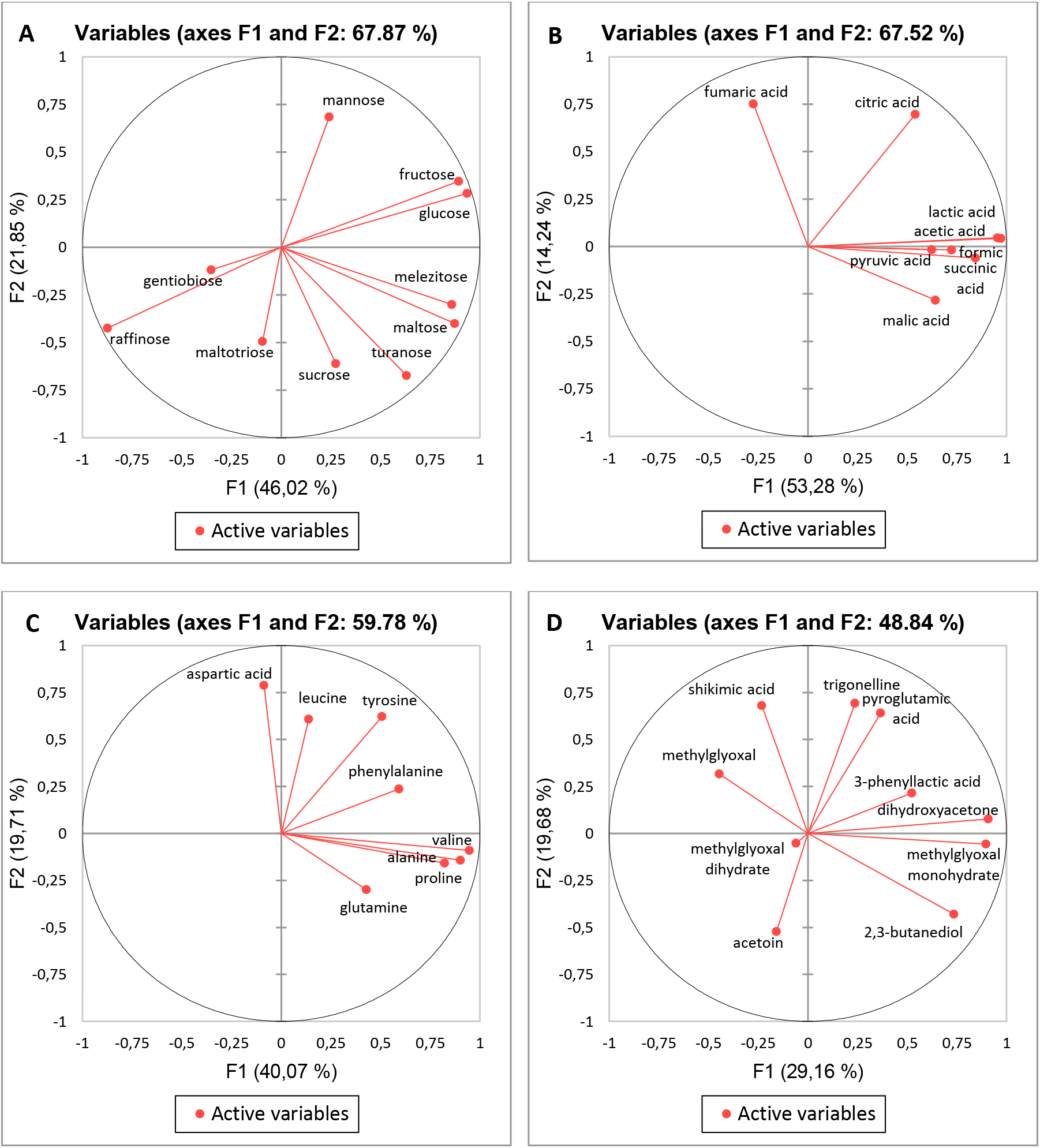
PCA loading plots for NMR metabolites: Sugars (**A**), Organic acids (**B**), Aminoacids (**C**), and Markers (**D**) in pot-honeys produced by *Geotrigona* (Geo), *Melipona* (Mel) and *Scaptotrigona* (Sca) Ecuadorian stingless bees.

One set of sugars is organized in the upper left portion (fructose, glucose, mannose), and two in the lower portion, left (gentiobiose, maltotriose, raffinose) and right (maltose, melezitose, sucrose, turanose) quadrant. (See Figure 8A) The first dimension F1 explains 46.02% variations. F1 explains 21.85% of the variation useful to position *Geotrigona* below and *Scaptotrigona* on the top. However separations of the three honey types are not complete. An interesting fact is that HMF and ethanol accounted for by 100% of the variance in another diagram not shown here, but these two parameters did not discriminate the honey types studied here. The organic acids were also distributed in three quadrants, in Figure 4B. Fumaric acid in the upper left, the major acids (acetic, citric and lactic) to the right, and formic, malic, pyruvic and succinic acids in the lower right. F1 explained 53.28% of the variance and F2 14.24%. A similar distribution for the aminoacids in Figure 4C. The aspartic acid is separated by the first dimension to the left from the remaining seven amino acids. Tyrosine, phenylalanine and leucine are in the upper right quadrant. Glutamine and a tight cluster of alanine, proline and valine are in the lower right quadrant. Looking at the aminoacids, the first and second dimensions accounted for by 59.78% of the variance –almost 10% less than sugars and organic acids. It was possible to see that *Geotrigona* was separated from *Melipona* by the first dimension, whereas *Scaptotrigona* was widespread in all space (not shown). Therefore, the PCA of amino acids did not discriminate the bee genus that originated the honey. In Figure 8D, the ten markers are distributed in all the space. Shikimic acid and methylglyoxal in the upper left, and acetoin and methylglyoxal dehydrate in the lower left. Trigonelline, pyroglutamic acid, 3-phenyllactic acid and dyhydroxyacetone in the upper right and in the lower right methylglyoxal monohydrate and the most abundant 2,3-butanediol.

The consensus configuration of the three Ecuadorian pot-honeys, shows their position in biplots of sugars and organic acids in Figure 9, with a similar resolution by sugars (9A) and organic acids (9B) to differentiate the bee genus of Ecuadorian honey. Both succeeded to cluster *Geotrigona* apart but *Melipona* and *Scaptotrigona* were not resolved. In the sugars biplot (Figure 9A), F1 separates *Geotrigona* to the left (gentiobiose, maltotriose and raffinose) from *Melipona* and *Scaptotrigona* to the right with the remaining seven sugars, F2 additionally positions *Melipona* in the upper quadrant (fructose, glucose, mannose) and *Scaptotrigona* in the lower position (maltose, melezitose, sucrose, turanose). In the organic acids biplot (Figure 9B), F1 separates *Melipona* and *Scaptotrigona* to the left with fumaric acid, and *Geotrigona* to the right with the other seven targeted NMR organic acids, but no further separation of *Melipona* and *Scaptotrigona* is achieved by F2. The first dimension explained 53.28% of the variance, and separated *Melipona* and *Scaptotrigona* honeys to the left, from *Geotrigona* to the right, where the major acids (acetic, citric and lactic) are. The second component (F2 14.24%) limits the position of *Melipona* in the lower quadrant but no further discrimination is achieved from *Scaptotrigona* honey types.

**Figure 9.**
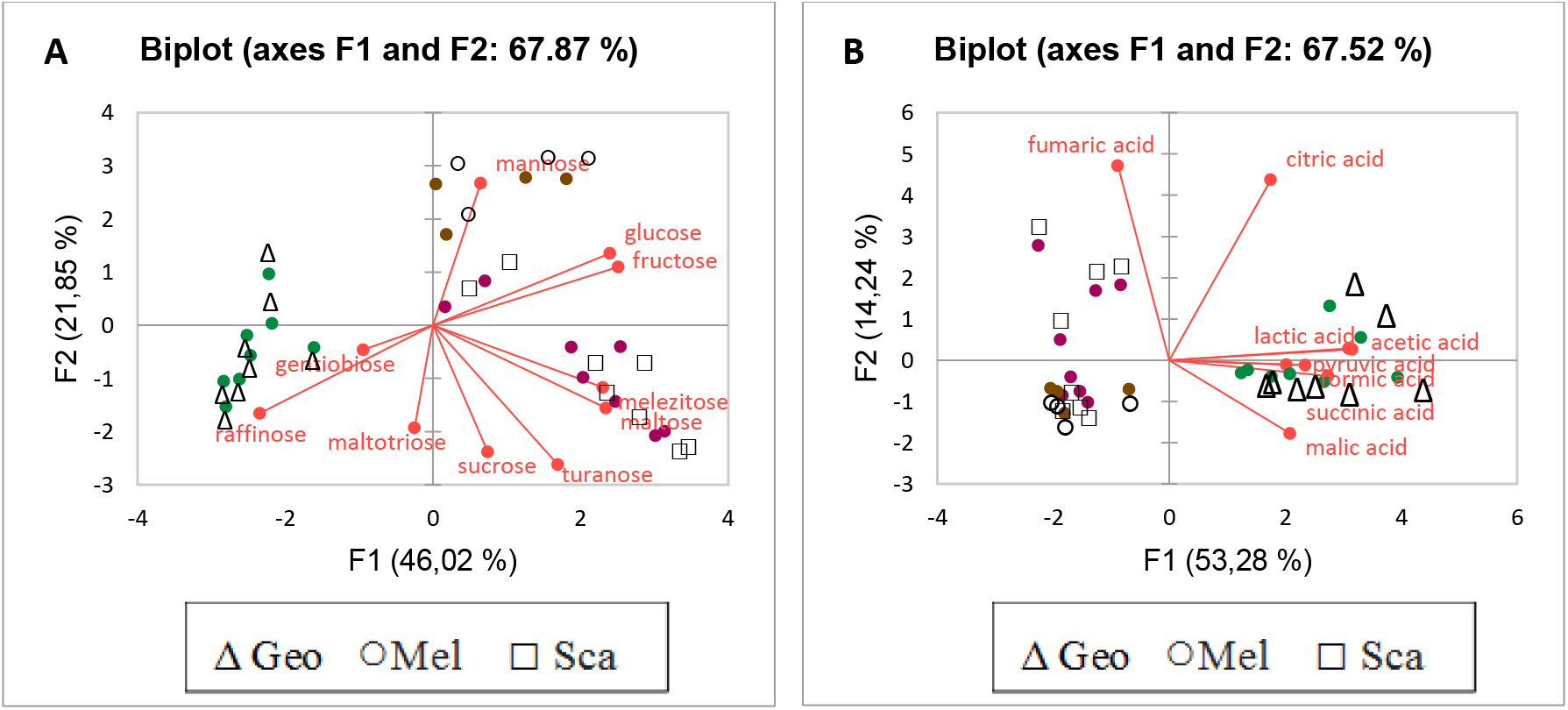
Biplots of all sugars **(A)**, and all organic acids **(B)** of pot-honeys produced by *Geotrigona* (Geo), *Melipona* (Mel) and *Scaptotrigona* (Sca) Ecuadorian stingless bees.

Hierarchical Cluster Analysis (HCA) performed to verify the grouping of the pot-honeys according to the ten sugars contents in Figure 9A formed two large clusters observed in the dendrogram, separating the *Geotrigona* (cluster 1) from the *Melipona* and *Scaptotrigona* (cluster 2) honeys. Similarly with the nine organic acid based dendrogram in Figure 2. This separation was not achieved with the two sets of eight amino acids and eight markers that are more related to the botanical origin. In the following section dendrograms will be compared with PCA.

The relationships between the targeted metabolites and the three bee genera studied here have a descriptive and quantitative value, but do not replace missing classic quality factors such as enzyme activity, moisture, ash content and electrical conductivity, besides sensory parameters like instrumental color. Indeed, *Melipona* and *Scaptotrigona* honey clusters were separated using DA in Venezuela (Vit et al., 1998) and PCA in Brazil (Avila et al., 2018), also the country of origin for *Geniotrigona* and *Heterotrigona* from Malaysia, and *Tetragonula* from Australia (Zawawi et al., 2022). Multiparametric analysis of these three collections of honey were not by NMR but included ash content for Venezuela and also mineral contents for the others. This fact points out the importance of inorganic elements and compounds to discriminate sets of honeys by multivariate analysis. Their origin in the nest is less studied than organic compounds. A reduction from ten to four variables was successful for DA separation of genus origin: Contents of nitrogen, reducing sugars, sucrose, and diastase activity. In contrast with the amino acids, not very discriminant in this work, total nitrogen was useful to separate *Melipona* and *Scaptotrigona* honeys from Venezuela (Vit et al., 1998). In targeted NMR, free acidity and pH are replaced by a set of organic acids, which besides the sugars were the two chemical groups able to explain 70% of the variance. Antioxidant and antibacterial activity are also used besides the elemental mineral contents to improve the multifactorial performance to discriminate botanical, geographical and entomological origin.

### 3.7 Assignment of stingless bee genus to three problem honeys

Three honeys received as “abeja de tierra” the ethnic name for underground stingless bees, “bermejo” the ethnic name for *Melipona mimetica*, and “cushillomishki” a Kichwa name, were tested by PCA and HCA of all sugars biplot in our database (Figure 6) to explore their entomological origin.

The PCA solution in Figure 10A positions each problem honey in each of the region of the three genera studied here. It is worth noting that being PCA a variable reduction approach based on correlations, inserting new honeys in the database affected the initial reference plot, as seen by comparing Figure 5A with 20 honeys and Figure 10A with 23, including the three honeys to be tested. Despite the allocated genus for each honey by PCA, in the HCA dendrogram with 20 reference pot-honeys + 3 problem samples (Figure 10B), the separation is not achieved. The three problem samples are located in the *Melipona-Scaptotrigona* cluster. This would mean the diagnostic for Ati is not a *Geotrigona* known as abeja de tierra. Ber and Cus are either *Melipona* or *Scaptotrigona*. The genus *Melipona* and the genus *Scaptotrigona* are too close for individual honey separations with the ten sugars or the organic acids measured in this work, that explained a 70% of the variance > 60 % by aminoacids > 50% by markers (See Figure 8). According to Figures 10A, Ati could be a *Geotrigona*, and Ber could be a *Melipona*. But in 10B Ati is not in the *Geotrigona* cluster. The multifactorial outcome needs further analysis for a diagnostic. Therefore, individual contents of sugars and organic acids from NMR profiles of these three honeys are given in Table 6.

**Figure 10.**
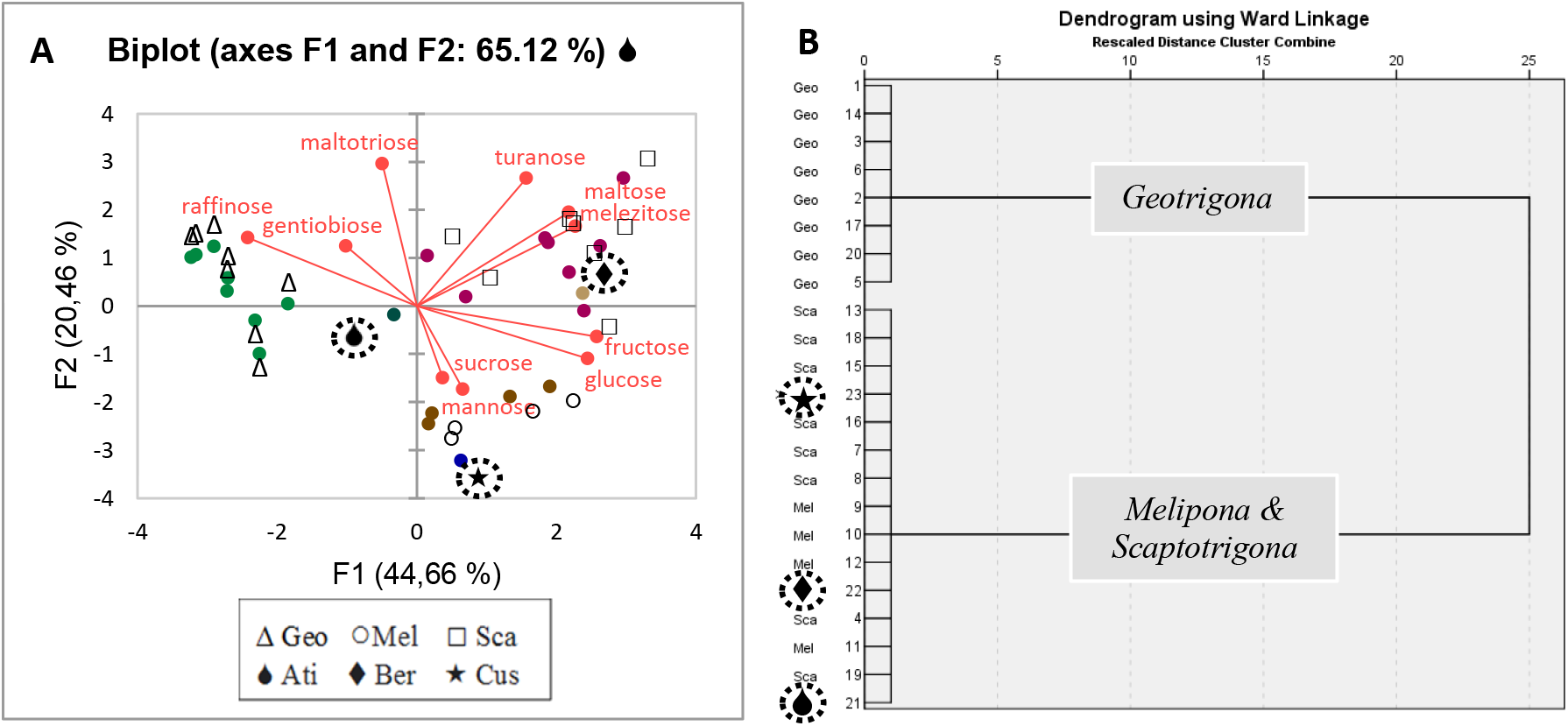
Biplot **(A)** and dendrogram **(B)** of all ten sugars in 23 honeys produced by *Geotrigona* (Geo), *Melipona* (Mel) and *Scaptotrigona* (Sca) Ecuadorian stingless bees, (see **Figs. 1B** and **5A** for 20 pot-honeys). In the sugars biplot, suggested solutions of the three additional problem honeys are highlighted with a dot-circle: 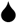**Ati** is in the left side of the biplot with Geotrigona, 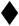**Ber** is not in the *Melipona* but in the *Scaptotrigona* region, positioned in the upper-right quadrant, and 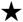**Cus** could be a *Melipona* honey positioned in the lower right quadrant. However in the dendrogram (6B) the three honeys were integrated in the *Melipona*-*Scaptotrigona* cluster. HCA by Euclidean distance using Ward’s method.

**Table 6.** Contents of sugars and organic acids (g/100 g honey) in three problem honeys.

| Honey | Sugars |  |  |  |  |  |  |  |  |  | Total |
| --- | --- | --- | --- | --- | --- | --- | --- | --- | --- | --- | --- |
|  | F | G | Ge | M | Ma | Me | Mt | R | S | T |  |
| Ati | 22.01 | 21.24 | 0.45 | 0.73 | 0.46 | 0.19 | 0.09 | 0.29 | 2.98 | 0.00 | 48.44 |
| Ber | 30.41 | 23.20 | 0.03 | 2.15 | 0.00 | 0.28 | 0.04 | 1.11 | 0.25 | 1.57 | 59.04 |
| Cus | 26.68 | 22.87 | 0.01 | 0.28 | 0.00 | 0.13 | 0.02 | 0.47 | 1.62 | 0.31 | 52.39 |
| Honey | Organic acids |  |  |  |  |  |  |  |  |  | Total |
|  | AcA | CitA | FoA | FuA | LacA | MalA | PygA | PyrA | QA | SucA |  |
| Ati | 0.65 | 0.07 | 0.05 | 0.00 | 0.93 | 0.01 | 0.01 | 0.00 | 0.00 | 0.00 | 1.72 |
| Ber | 0.17 | 0.00 | 0.00 | 0.00 | 0.17 | 0.00 | 0.00 | 0.00 | 0.00 | 0.00 | 0.34 |
| Cus | 0.21 | 0.02 | 0.00 | 0.00 | 0.23 | 0.00 | 0.00 | 0.00 | 0.00 | 0.00 | 0.46 |

**Honey:** **Ati** Abeja de tierra *Geotrigona*, **Ber** Bermejo *Melipona*, **Cus** Cushillomishki Kichwa name. **Sugars:** **F** fructose, **G** glucose, **Ge** Gentiobiose, **M** maltose, **Mn** mannose, **Me** melezitose, **Mt** maltotriose, **R** raffinose, **S** sucrose, **T** turanose. **Organic acids:** **AcA** acetic acid, **CitA** citric acid, **FoA** formic acid, **FuA** fumaric acid, **LacA** lactic acid, **MalA** malic acid, **PygA** pyroglutamic acid, **PyrA** pyruvic acid, **QA** quinic acid, **Suca** succinic acid.

From Table 6, the F+G of Ati 43.25 g/100g is too high for a *Geotrigona* honey, see the range [13.08 - 24.50] in Table 1, and the total organic acids are too low (1.72) compared to Table 3a (4.44). Ber has 53.61g/100g F+G, which is within the range of *Melipona* and *Scaptotrigona* [49.12 - 56.10], as well as 0.17 for each acetic acid and lactic acid lower than averages in Table 3 [1.2-1.6]. The maltose content (2.15) is in the *Scaptotrigona* range [0.92-2.75], higher than Geo and Mel. Except for the lower maltose (0.28) content, Cus is similar to Mel, with lower F+G (49.55), slightly higher acetic acid (0.21) and lactic acid (0.23). Therefore, based in this descriptive analysis, Cus is more likely a *Melipona* honey, Ber could be a *Scaptotrigona* or a *Melipona* sp. with higher maltose content and if Ati is not a *Geotrigona*, sugars were possibly added to that honey, not cane sugar but another honey perhaps from *Apis mellifera*, able to increase fructose and glucose contents. The F+G content in Ati is too high for *Geotrigona*, and too low for *Melipona* or *Scaptotrigona*. The organic acids, particularly, the acetic acid and the lactic acid are too low for *Geotrigona* and too high for *Melipona* and *Scaptotrigona*. Also, if Ati is not a *Geotrigona* honey, could be from another genus not studied here.

### 3.8 The Honey Authenticity Test by Interphase Emulsion (HATIE) revealed pot-honey biosurfactants in Scaptotrigona sp

A diagram in Figure 11 shows two patterns for genuine pot-honey: 1. One phase for *Scaptotrigona* and three phases for *Geotrigona* and *Melipona*. The HATIE authenticity test used with genuine pot-honeys becomes a Honey Biosurfactant Test (HBT). *Scaptotrigona* honey has a suspected microbial origin surfactant causing that distinctive pattern that makes it an innovative entomological origin fingerprint for this genus, compared to *Geotrigona* and *Melipona*, in this set of pot-honeys. There are 22 species of *Scaptotrigona* to be tested.

**Figure 11.**
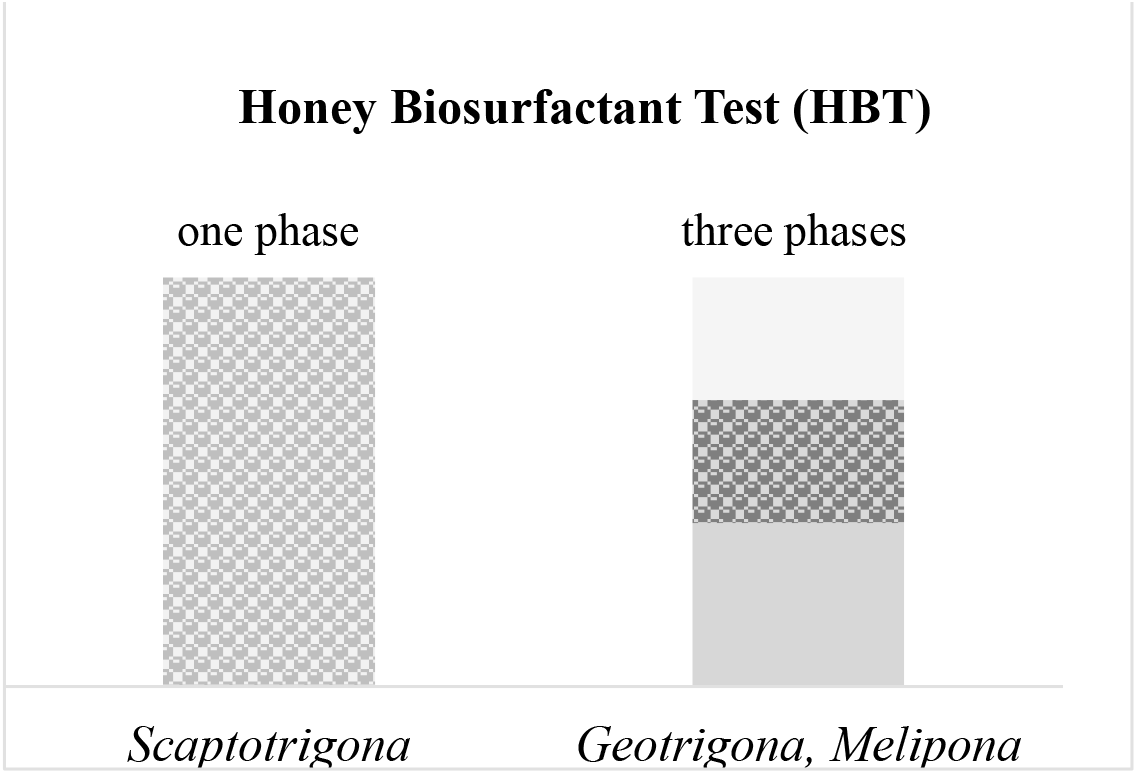
Diagram for genuine Honey Biosurfactant Test

### 3.9 Searching stingless bee markers in pot-honey

Bee-related lipophilic signals in the central region of honey NMR spectra (3.5-4.5 ppm), are more likely to be entomological markers (Vit et al., 2015). However, considering the selectivity of food choices in different habitats, diverse compound classes (amino acids, organic acids and sugars) are showing a functional role for entomological assignment of pot-honey. They would ideally indicate what kind of bee produced the honey –at least comparing bees from the same region. Their presence and concentrations represent different origins and functions. An unusual HPLC high maltose content was proposed as a predictor of entomological origin for Venezuelan pot-honeys using multivariate HCA, a cluster of *Melipona*-*Scaptotrigona* was separated from *Frieseomelitta*-*Nannotrigona*-*Tetragonisca* (Vit et al., 1998a). A further discriminant analysis (DA) with ten quality factors separated *Melipona*, *Scaptotrigona* and other Trigonini into three clusters (Vit et al., 1998b). Phylogenetically this pattern is interesting given that *Melipona* and *Scaptotrigona* are not closely related (Roubik, 1992; Rasmussen and Cameron, 2010). The maltose source is a minor floral nectar, more related to transglycosilation enzymes added by bees to make honey (Low et al., 1988). An NMR-LCMS-based metabolomics approach of stingless bee honey from Malaysia suggested discriminant metabolites were one amino acid, two organic acids and four sugars: These were D-fructofuranose in honey from *Heterotrigona itama*, α-D-glucose, β-D-glucose, D-xylose from *Geniotrigona thoracica* and L-lactic acid, acetic acid, L-alanine from *Tetrigona apicalis* (Razali et al., 2018). In our data, raffinose was a sugar that discriminated *Geotrigona* (5.03 g/100 g) from a *Melipona* and *Scaptotrigona* mixed cluster. (See Figure 1B). However, raffinose also was in the sugar spectra (0.01-0.17 g/100 g) of *A. mellifera* honey from Korea (Jang et al., 2016), (0.03-0.08 g/100 g) from Spain (Pascual-Matéa et al., 2018), and (0.65 g/100 g) of *M. ferruginea* from Tanzania (Popova et al., 2021). Therefore, it is not an entomological marker specific for *Geotrigona*, but it was a discriminant sugar compared to *Melipona* and *Scaptotrigona* honeys from Ecuador in this study. A genetic bee-origin of chemical profiles in honey has not been approached as the vast literature on geographic and plant origin –bee diet– It is referred to as entomological origin mainly for comparative scopes of compositional (chemical, botanical, physical) and functional (antibacterial, anticancer, anti-inflammatory, antioxidant, cytotoxic, sensory) properties. This possible influence of stingless bee genus would help our understanding of environmental or other factors, if in fact honey production has a component driven by bee phylogeny or taxonomic group. Navigating tricky questions regarding stingless bee phylogeny is an expression used by Grüter (2020) in the preface of his book, to anticipate the complexity of the topic.

Bees are known to be selective in their choices of pollen and nectar sources. *Geotrigona acapulconis* collects nectar from ‘Hass’ cultivars of avocado *Persea americana* in its native Mexico. Among the *Geotrigona argentina* floral resources studied in 146 pollen pots and combined pot-honey from four nests, major nectar sources overlapped major pollen sources (Vossler et al., 2010). *Tetragonula carbonaria* and *Austroplebeia australis* use of sympatric resources may have phylogeographic origins to avoid or reduce competition: 1. *T. carbonaria* higher flight activity enabled faster sugar intake from a broader plant spectrum than *A. australis*, 2. The smaller *A. australis* colonies collected higher sugar concentrations from a narrower plant spectrum (Leonhardt et al., 2014a). Accordingly, these distinctive behavioral traits could convey the chemical composition of honey. Thus, studying quality traits in honey seems possible.

For example, after feeding sucrose solutions to *T. carbonaria* a fake honey with (64-72) g trehalulose/100 g honey, (18-23) erlose, (9-12) fructose, and minor glucose was produced in contrast to a fructose-glucose (25:25:50W) feeding that produced a 58 g/100 g total sugars were the F:G ratio fed remained unchanged. These authors hypothesized a major trehalulose honey from stingless bees feeding high sucrose floral nectar (Hungerford et al., 2021). A feral small stingless bee *Frieseomelitta* aff. *varia* from Venezuela, also had a surprisingly low 6.9 g glucose/100 g honey, possibly the very high (56.9) maltose-like HPLC sugar was trehalulose (Vit et al., 1998a). Being trehalulose a rare sucrose isomer α-(1→ 1) glucose-fructose, bee-nest-enzymes involved in its anabolic/catabolic pathways could be a potential candidate for a valid long-term physiological trait, in agreement with behavioral observations by Leonhardt et al. (2014a). Nectars colonized by microbial species affect bee foraging behavior based on chemical changes they cause to the nectar (Good et al., 2014), nectar volatiles influence food preference and acceptability (Rering et al., 2018), pollen and yeast-based emissions were compared (Rering et al., 2020). A floral nectar colonizing yeast *Metschnikowia* that may reach the bee nest, was a common genus in honey bee guts (Kakumanu et al., 2016). Genes codifying for α-glucosidases from the nectar-yeast *M. gruessii* and *M. reukaufii* were expressed (Garcia-Gonzalez, 2019), and produced rare sugars present in honey (erlose, isomelezitose, trehalulose) after sucrose splitting activity (Garcia-Gonzalez, 2020).

A note on phylogenetic traits. They are diverse features essential to understanding the evolution of biodiversity, connecting microevolutionary processes of natural selection to macroevolutionary patterns of phenotypic evolution (Hansen and Martins, 1996). Not only morphological traits, but also long-term changes in physiology and behavior are traits of phylogeny –sharing features. A phylogenetic or family tree is a graphic indicator with branches (relationships) used to depict a history of evolution for a group of taxa (i.e., species, genus). The term “phylogeny” was coined in German by Ernst Haeckel in 1870 (the word is of Greek derivations where phylos φῦλον = race + geneia γενεᾱ’ = birth or origin). Dendrograms of stingless bee taxa related by the chemical profiles of their honey can inform us on the skills to: 1. Select sugary materials available from the environment, 2. Apply bee-physiology for transformations in the nest, i.e. bee-enzymes 3. Select associated microbiota for further processing, defense, and preservation of a moist sugary substrate from spoilage in the warm nest. Phylogenetic and geographical signal of host-specific LAB in *Tetragonula* and *Austroplebeia* stingless bees contributed to the evolutionary history of the bee-LAB association (Leonhardt et al., 2014b).

In a morphological trait such as the cuticular hydrocarbon profile, bee species may be distinctive: 1.Different drivers can affect the chemical profile based on functionality, 2. The ability to acquire environmental materials (chemical compounds) is correlated strongly with functionality, as well as the overall phylogenetic placement of the bee (Leonhardt et al., 2013). What physiological traits instead of a morphological trait would convey a similar correlation for honey? A multitask behavior to preserve the quality of honey processing for the health of bee colonies (Leonhardt, 2017). The nutritional choices made by stingless bees on food resources, may additionally become a solid source of information for long-term conservation plans (Leonhardt et al., 2020). Furthermore, a bee-derived marker, was that of associated microbiota-derived marker. More microbial-origin markers of the stingless bee nest are waiting to be discovered in the future, in addition to the overwhelming quantities of acetic acid and lactic acid they produce. A *Bacillus* marker or a related AAB-LAB, a yeast or fungal fingerprints? Arabitol and mannitol of cerumen extracts from stingless bee nests are sweet sugar alcohols (polyols) that may be of interest as fungal origin (Popova et al., 2021). Another answer now is a suspected *Starmerella* yeast associated with *Scaptotrigona* sp., one possibility is *Starmerella bombicola*, known for the extracellular sophorolipid biosynthesis with surfactant action. This fingerprint was not detected by ^1^H-NMR but by the HATIE, honey authenticity test based on interphase emulsion formed after shaking a honey water dilution with diethyl ether.

## 4. Conclusions

From the 41 parameters studied here by targeted ^1^H NMR, the amino acid isoleucine, the organic acid quinic acid, and the marker kynurenic acid were not detected in the Ecuadorian honeys. Therefore, the multivariate analysis was conducted for 10 sugars, 9 organic acids, 9 amino acids and 8 markers. Sugars and organic acids were more discriminant of the bee genus than amino acids and markers. A test of three problem honeys revealed multifactorial analysis limitations and the need of descriptive parameters for a preliminary fit of these honeys according to their bee-genus, with the incipient 20-honey database presented here. A solid database needs to grow for useful comparisons with similar honeys and also to know the exceptions.

The targeted NMR chemical profiles are useful for a multi-purpose reference database to check an unknown pot-honey or suspected of adulteration, mislabeled or fake. Such a database is growing for botanical, entomological and geographical origins, and adulteration sources from various countries. With so many floral and entomological resources, the biodiversity of honeys in Ecuador is a treasure to be discovered. Our results with limited number of honeys deserve continuation, with few honey quality control data besides the NMR. Considering that the pot-honey composition bears a relationship of taxa from three kingdoms represented by plant-bee-microbiota, facts not explained by the botanical and entomological origin, may have a microbiota answer in the nest. The discovery of biosurfactant activity in *Scaptotrigona* honey evidenced a microbial origin that needs identification as well as the chemical nature of the biosurfactant.

The scientific interest to investigate pot-honey composition is a slow cumulative effort to contribute to meliponine scientific research community for understanding the bee processes and for regulatory purposes, which eventually will support stingless bee keeping, agriculture, and consumers. A norm for the Ecuadorian honey produced by stingless bees is needed and the NMR data reported here is a support to establish required honey quality standards.

The term foodomics coined by Alejandro Cifuentes (2009) as “A discipline that studies the Food and Nutrition domains through the application of omics technologies” is needed for pot-honey research and discoveries by specialized teams on genomics, transcriptomics and proteomics, besides the metabolomics that has already started. The consistent influence of stingless bee genus in honey would help understanding environmental or other factors, if in fact stingless bee honey processing has a component driven by bee phylogeny and associations with microbes having multifaceted functions including nutritional traits. The biosurfactant activity of *Scaptotrigona* honey revealed by the test suggested microbial associations originating that visual behavior. Our conclusion is for this set of Ecuadorian pot-honeys. Echeverrigaray et al. (2021) found *Starmerella bombicola* in the Brazilian honey of two *Scaptotrigona* spp., *S. bipunctata* and *S. ederi* but not in *Scaptotrigona tubiba*. Therefore, the microbial association with stingless bees is species specific for *Scaptotrigona*, not genus specific as in our Ecuadorian study with only one species of *Scaptotrigona* sp.

## Supporting information

Table S1 Targeted NMR

## CRediT authorship contribution statement

**Patricia Vit**: Conceptualization, Field work, Database, Analyzed and interpreted the data - original draft, Writing - review & editing. **Jane van der Meulen**: Formal analysis, Database, **Silvia RM Pedro**: Stingless bee genera, Database. **Isabelle Esperança**: Analyzed and interpreted the data, review & editing. **Rahimah Zakaria**: Analyzed the data, review & editing. **Gudrun Beck**: Resources, **Favian Maza**: Resources, Prometeo supervisor.

## Acknowledgements

To stingless bee keepers from Ecuador. David W. Roubik from Smithsonian Tropical Research Institute in Ancon, Panama, and Michael S. Engel from the Division of Entomology, Natural History Museum, and Department of Ecology & Evolutionary Biology, University of Kansas, Lawrence kindly edited the intricated final section. Martin Linkogel from Quality Services International GmbH, Bremen, Germany, for his professionalism to facilitate current institutional contacts. To Omar Malagón from Universidad Técnica Particular de Loja, for the map of Ecuador. To anonymous reviewers who improved our expressions in the manuscript. To CDCHTA from Universidad de Los Andes, Mérida, Venezuela. To Universitá Politecnica delle Marche, Ancona, Italy.

