## Supplementary material for "Impact of genus (*Geotrigona*, *Melipona*, *Scaptotrigona*) in the ^1^H-NMR organic profile and authenticity test of honey processed in cerumen pots by stingless bees in Ecuador": Table S1 Targeted NMR

**Table S1.** Targeted 41 organic metabolites, molecular formula, chemical structure, regions of the $^{1}$H-NMR spectra (ppm), signal type and $^{1}$H-NMR spectra.

| No. | **Organic metabolite** Molecular formula | **Chemical structure**[1] | **$^{1}$H-NMR range (ppm)**[2] | **Signal type**[2] | **$^{1}$H-NMR spectra**[1] |
|---|---|---|---|---|---|
| | **Sugars** | | | | |
| 1 | Fructose<br>$C_6H_{12}O_6$ | 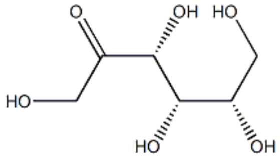 | 4.12-4.07 | doublet | - |
| 2 | Glucose<br>$C_6H_{12}O_6$ | 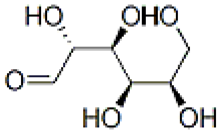 | 4.65-4.62 | singlet | 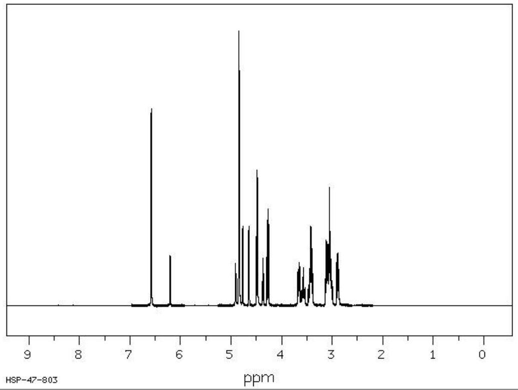 |
| 3 | Sucrose<br>$C_{12}H_{22}O_{11}$ | 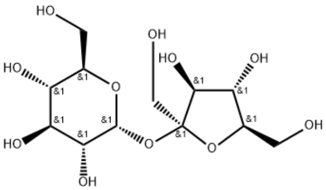 | 4.22-4.19 | doublet | 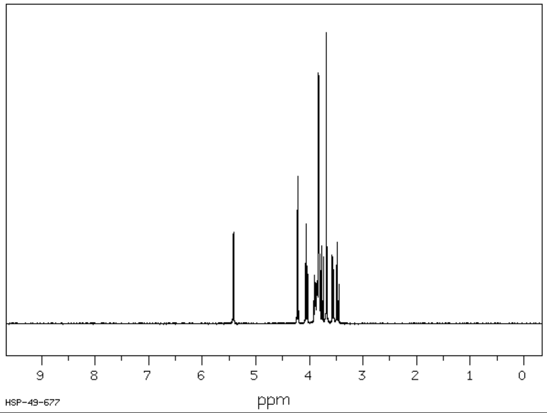 |

| | | | | | |
|---|---|---|---|---|---|
| 4 | Gentiobiose<br>$C_{12}H_{22}O_{11}$ | 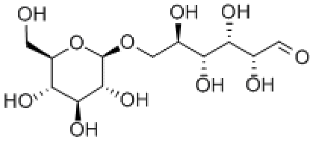 | 4.51-4.48 | doublet | 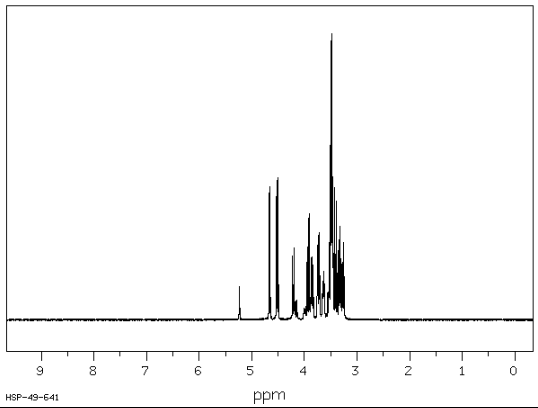 |
| 5 | Maltose<br>$C_{12}H_{22}O_{11}$ | 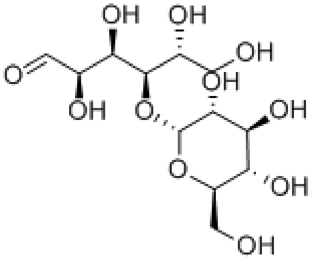 | 5.18-5.17 | doublet | - |
| 6 | Maltotriose<br>$C_{18}H_{32}O_{16}$ | 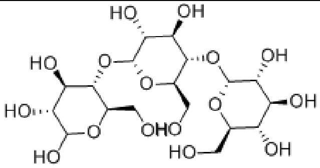 | 3.29-3.26 | doublet | - |
| 7 | Mannose<br>$C_6H_{12}O_6$ | 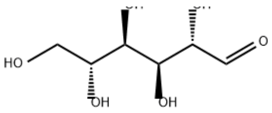 | 5.39-5.38 | doublet | 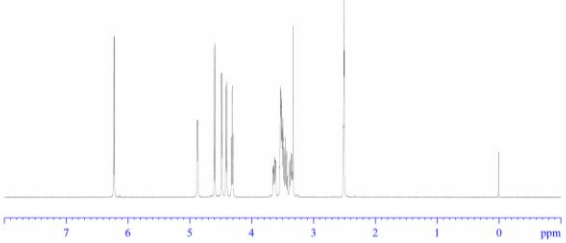 |

| | | | | | |
|---|---|---|---|---|---|
| 8 | Melezitose<br>$C_{18}H_{32}O_{16}$ | 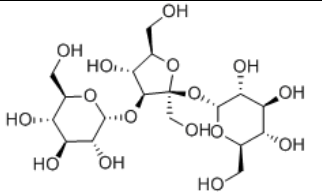 | 5.46-5.42<br>4.29-4.27 | doublet | - |
| 9 | Raffinose<br>$C_{18}H_{32}O_{16}$ | 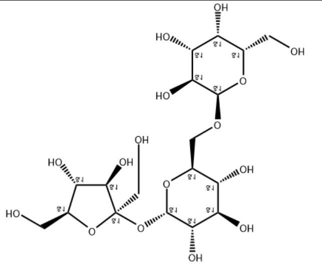 | 4.99-4.97 | doublet | - |
| 10 | Turanose<br>$C_{12}H_{22}O_{11}$ | 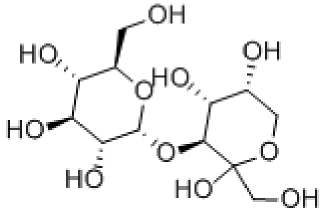 | 5.3-5.27 (d) | doublet | - |
| | **Indicator of quality** | | | | |
| 11 | **5-Hydroxymethylfurfural**<br>$C_6H_6O_3$ | 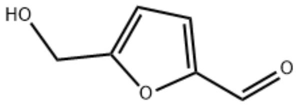 | 9.46-9.43 | singlet | 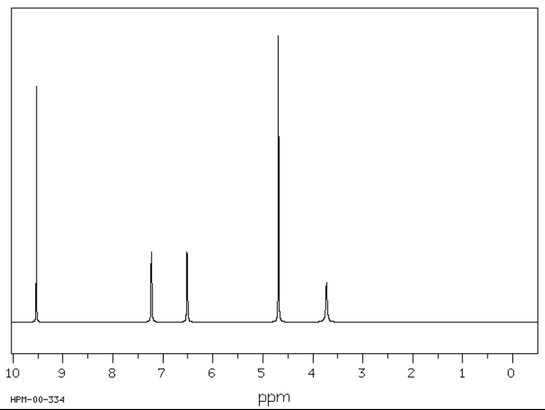 |
| | **Alcohols** | | | | |

| | | | | | |
|---|---|---|---|---|---|
| 12 | **Ethanol**<br>$C_2H_6O$ | 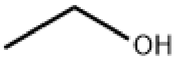 | 1.14-1.20 | triplet | 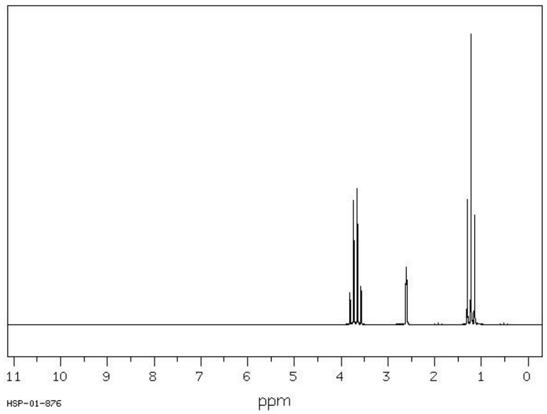 |
| | **Aliphatic Organic Acids (AOA)** | | | | |
| 13 | Acetic Acid<br>$CH_3COOH$ | 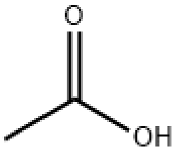 | 2.09-2.08 | singlet | 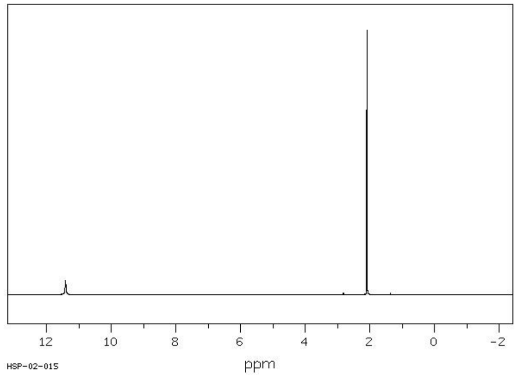 |
| 14 | Citric Acid<br>$C_6H_8O_7$ | 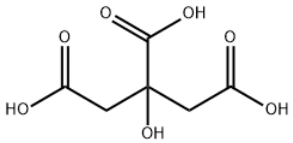 | 2.99-2.92<br>2.84-2.77 | singlet | 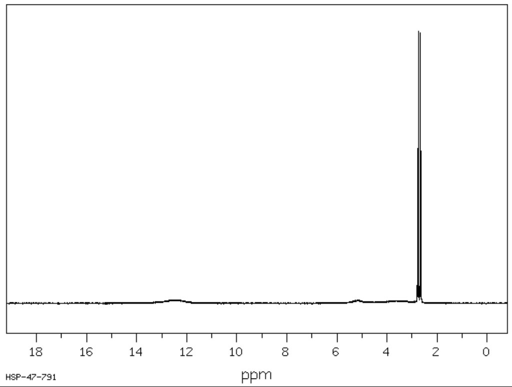 |

| | | | | | |
|---|---|---|---|---|---|
| 15 | Formic Acid<br>$CH_2O_2$ | 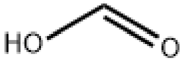 | 8.31-8.23 | singlet | 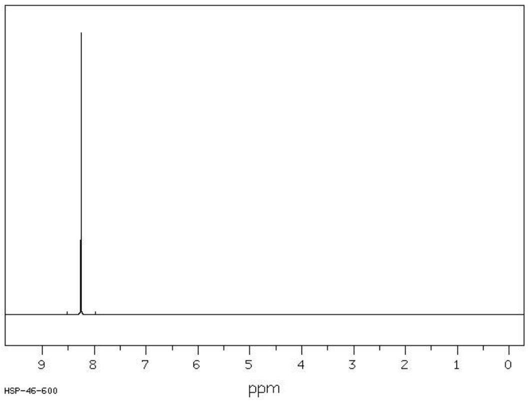 |
| 16 | Fumaric Acid<br>$C_4H_4O_4$ | 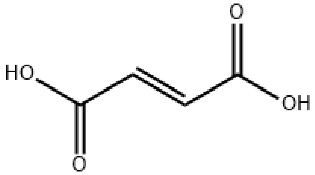 | 6.75-6.72 | singlet | 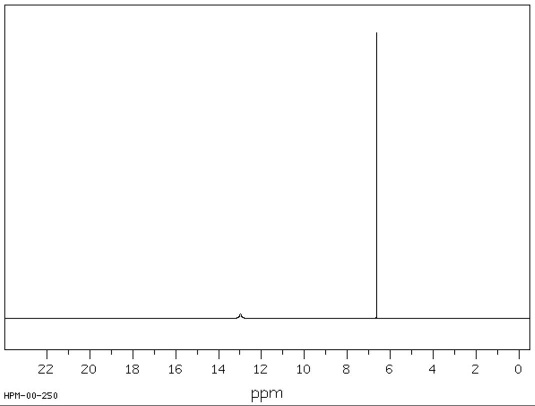 |
| 17 | Lactic Acid<br>$C_3H_6O_3$ | 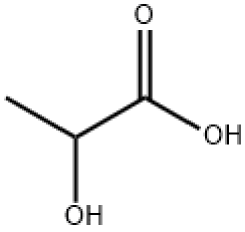 | 1.38-1.42 | doublet | 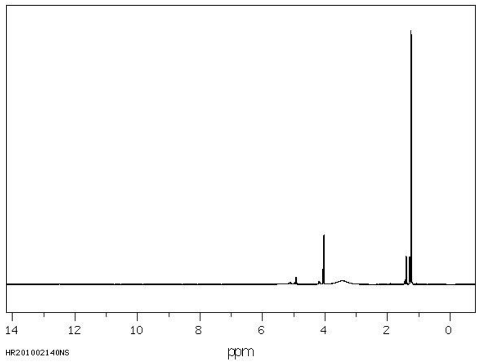 |

| | | | | | |
|---|---|---|---|---|---|
| 18 | Malic Acid<br>$C_4H_6O_5$ | 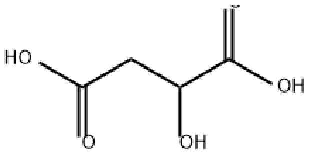 | 2.93-2.85<br>2.82-2.74 | double doublet | 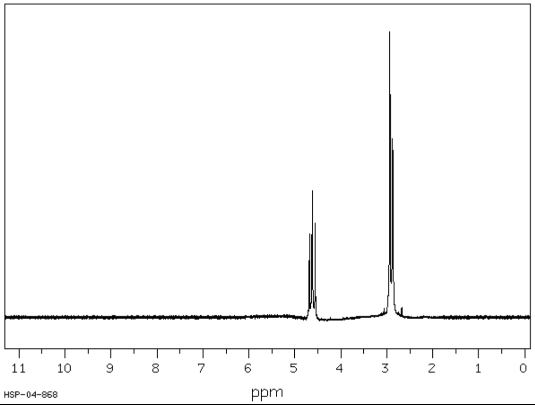 |
| 19 | Pyruvic Acid<br>$C_3H_4O_3$ | 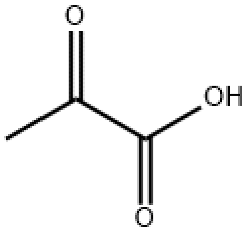 | 2.37-2.37 | singlet | 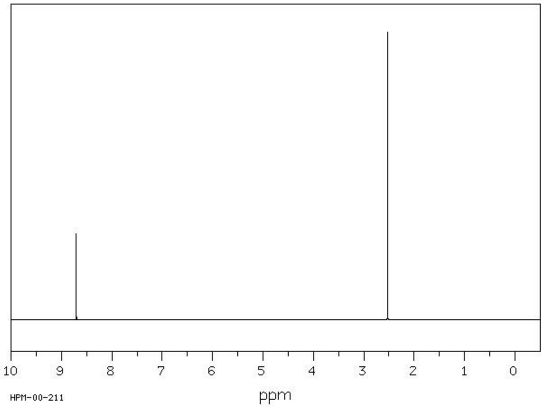 |
| 20 | Quinic Acid<br>$C_7H_{12}O_6$ | 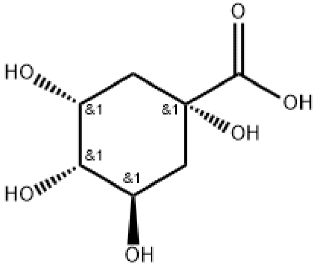 | 1.95-1.85 | double doublet | 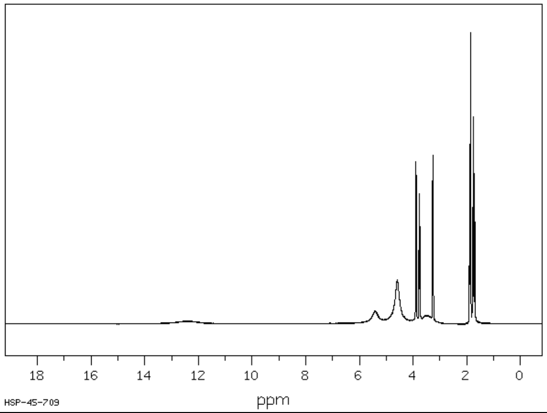 |

| | | | | | |
|---|---|---|---|---|---|
| 21 | Shikimic Acid<br>$C_7H_{10}O_5$ | 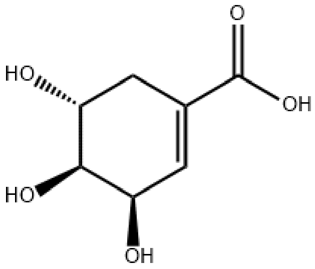  | 6.82-6.78 | multiplet | 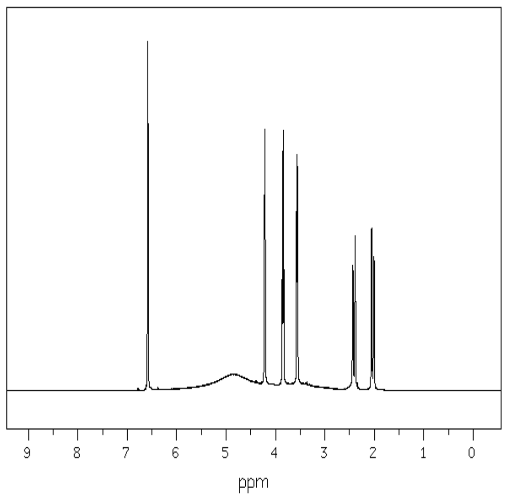  |
| 22 | Succinic Acid<br>$C_4H_6O_4$ | 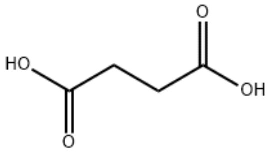  | 2.67-2.64 | singlet | 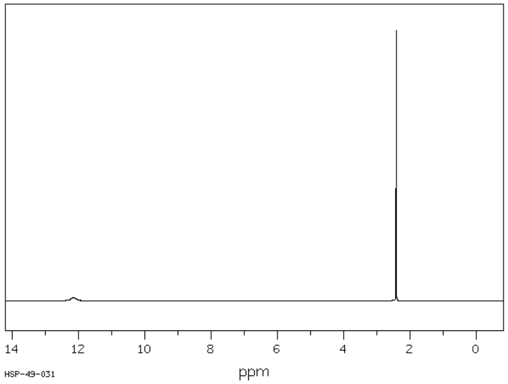  |
| | **Amino Acids** | | | | |

| | | | | | |
|---|---|---|---|---|---|
| 23 | Alanine<br>$C_3H_7NO_2$ | 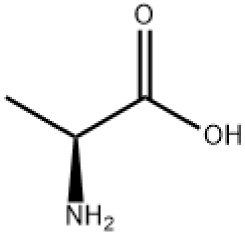 | 1.50-1.47 | doublet | 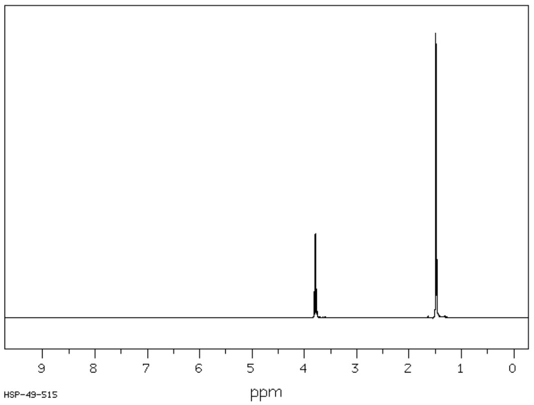 |
| 24 | Aspartic acid<br>$C_4H_7NO_4$ | 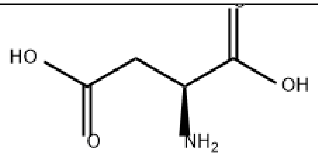 | 2.95-2.91 | doublet | |
| 25 | Glutamine<br>$C_5H_{10}N_2O_3$ | 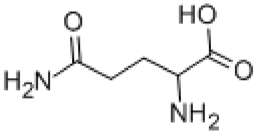 | 2.48-2.42 | multiplet | 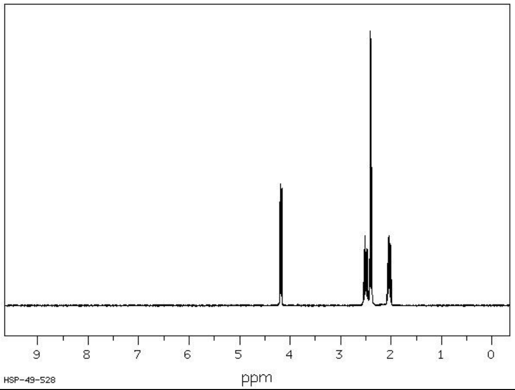 |
| 26 | L-Isoleucine<br>$C_6H_{13}NO_2$ | 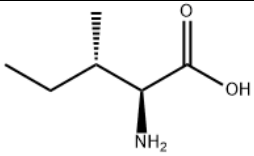 | 1.00-0.99<br>0.93-9.92 | multiplet | - |

| | | | | | |
|---|---|---|---|---|---|
| 27 | Leucine<br>$C_6H_{13}NO_2$ | 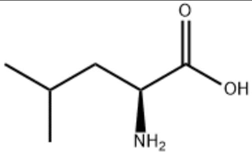  | 0.96-0.95<br>0.94-0.93 | multiplet | - |
| 28 | Phenylalanine<br>$C_9H_{11}NO_2$ | 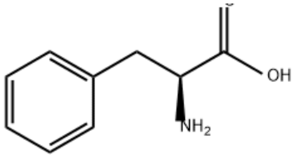  | 7.39-7.45 | multiplet | 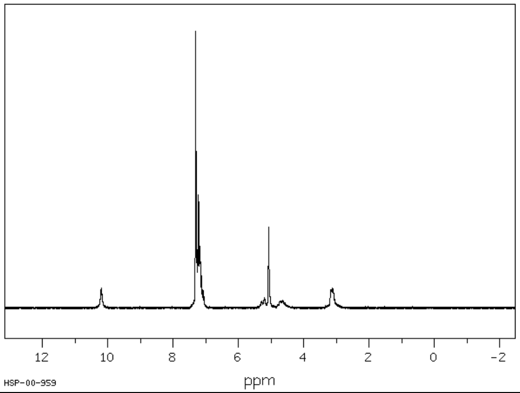  |
| 29 | Proline<br>$C_5H_9NO_2$ | 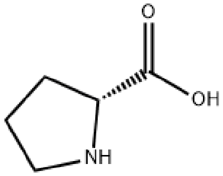  | 2.40-2.29<br>2.11-2.07 | multiplet | 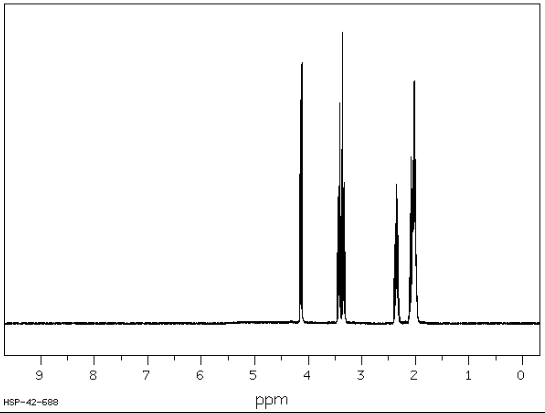  |

| | | | | | |
|---|---|---|---|---|---|
| 30 | Pyroglutamic acid<br>$C_5H_7NO_3$ | 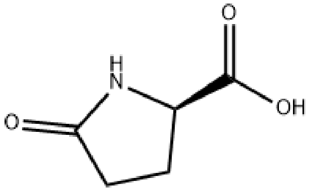 | 2.43-2.39 | | 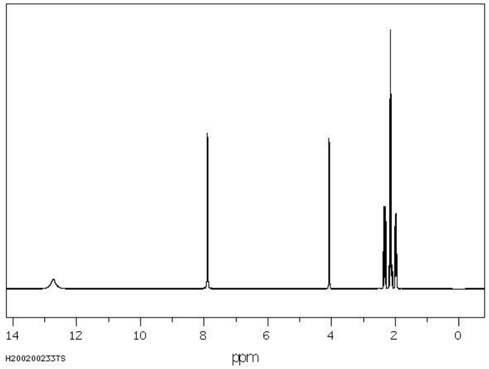 |
| 31 | Tyrosine<br>$C_9H_{11}NO_3$ | 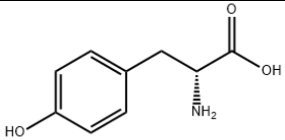 | 6.92-6.86 | multiplet | - |
| 32 | Valine<br>$C_5H_{11}NO_2$ | 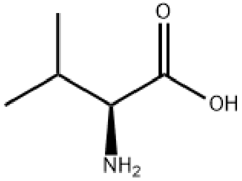 | 1.05-1.02<br>0.99-0.97 | doublet | - |
| | **Botanical markers** | | | | |
| 33 | Acetoin | 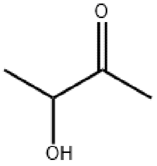 | 2.23-2.22<br>1.38-1.35 | singlet<br>doublet | 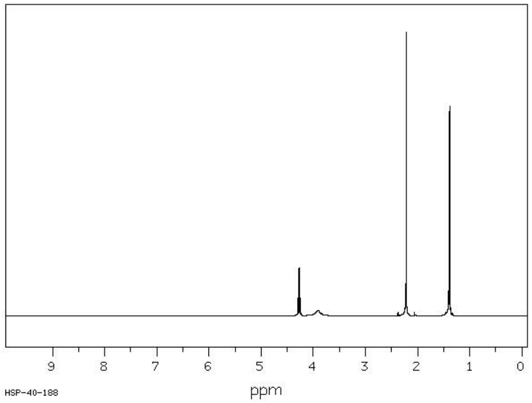 |

| | | | | | |
|---|---|---|---|---|---|
| 34 | 2,3-Butanediol | 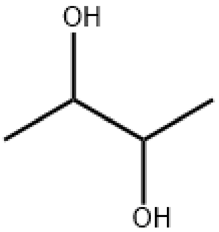  | 1.14-1.12 | doublet | 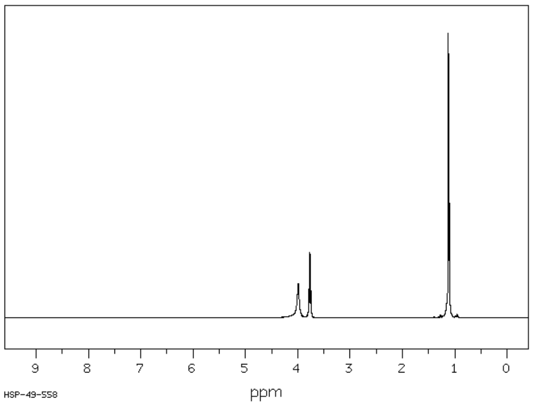  |
| 35 | Dihydroxyacetone | 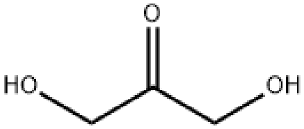  | 4.41-4.40 | singlet | |
| 36 | Kynurenic acid | 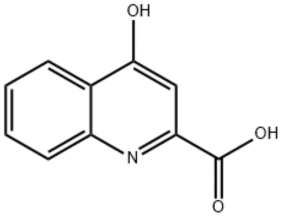  | 7.08-6.98 | multiplet | 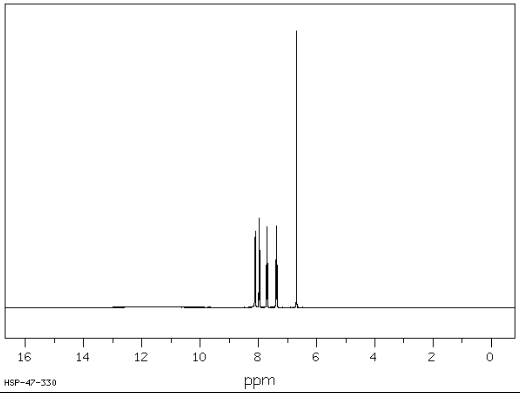  |
| 37 | Methylglyoxal $C_3H_4O_2$ | 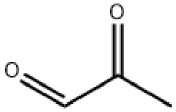  | 1.99 | singlet | . |
| 38 | Methylglyoxal dihydrate $C_3H_7O_4$ | - | 1.37-1.36 | singlet | - |

| | | | | | |
|---|---|---|---|---|---|
| 39 | Methylglyoxal monohydrate $C_3H_6O_3$ | 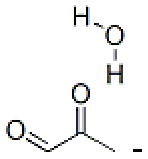 | 2.29-2.28 | singlet | - |
| 40 | 3-Phenyllactic acid $C_9H_{10}O_3$ | 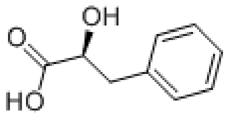 | 3-20.940 | double doublet | 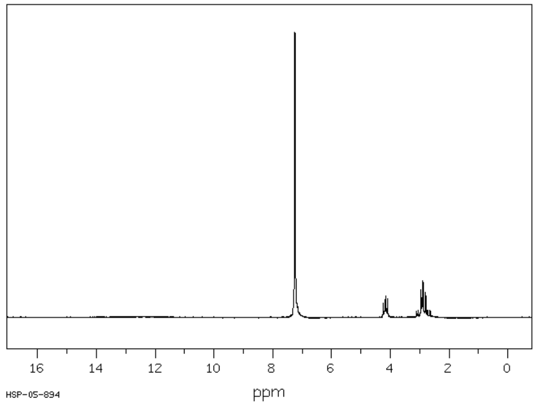 |
| 41 | Trigonelline $C_7H_7NO_2$ | 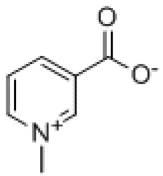 | 9.17-9.10 | triplet | 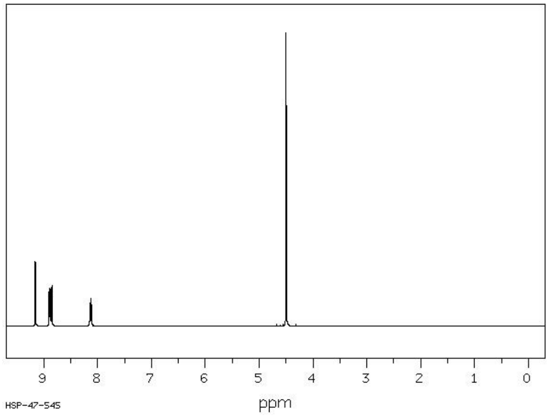 |

[1]Chemical structures and $^{1}$H-NMR spectra from Chemical Book https://www.chemicalbook.com/SpectrumEN
[2]M. Díaz QSI Bremen, Germany 2022
